# A platform for dissecting force sensitivity and multivalency in actin networks

**DOI:** 10.1101/2023.08.15.553463

**Authors:** Joseph T. Levin, Ariel Pan, Michael T. Barrett, Gregory M. Alushin

## Abstract

The physical structure and dynamics of cells are supported by micron-scale actin networks with diverse geometries, protein compositions, and mechanical properties. These networks are composed of actin filaments and numerous actin binding proteins (ABPs), many of which engage multiple filaments simultaneously to crosslink them into specific functional architectures. Mechanical force has been shown to modulate the interactions between several ABPs and individual actin filaments, but it is unclear how this phenomenon contributes to the emergent force-responsive functional dynamics of actin networks. Here, we engineer filament linker complexes and combine them with photo-micropatterning of myosin motor proteins to produce an *in vitro* reconstitution platform for examining how force impacts the behavior of ABPs within multi-filament assemblies. Our system enables the monitoring of dozens of actin networks with varying architectures simultaneously using total internal reflection fluorescence microscopy, facilitating detailed dissection of the interplay between force-modulated ABP binding and network geometry. We apply our system to study a dimeric form of the critical cell-cell adhesion protein α-catenin, a model force-sensitive ABP. We find that myosin forces increase α-catenin’s engagement of small filament bundles embedded within networks. This activity is absent in a force-sensing deficient mutant, whose binding scales linearly with bundle size in both the presence and absence of force. These data are consistent with filaments in smaller bundles bearing greater per-filament loads that enhance α-catenin binding, a mechanism that could equalize α-catenin’s distribution across actin-myosin networks of varying sizes in cells to regularize their stability and composition.

## Main

The actin cytoskeleton mediates mechanical interactions between cells and their local environments, facilitating cell movement and the integration of cells into tissues (1). The cytoskeleton’s diverse functions are implemented by subcellular networks composed of actin filaments (F-actin) and more than 100 different actin-binding proteins (ABPs) (2). The geometric arrangement of actin filaments within these networks (i.e. relative filament orientations, inter-filament angles, and spacing), as well as their local ABP composition, determines their mechanical properties and biochemical activities (3). While signaling processes that converge on Rho GTPases to initiate the formation of specific classes of actin networks are well-established (4, 5), our understanding of feedback mechanisms which dynamically tune the geometry and composition of networks as they engage in mechanical functions remains limited.

Multiple actin networks, featuring divergent architectures optimized for specific functions, frequently co-exist within the same cytoplasm. Branched actin networks, defined by the presence and activity of the ARP2/3 complex, polymerize at the leading edge of migrating cells with characteristic 70° branching angles to push the cell membrane forward and drive migration (6, 7). Contractile co-linear actin bundles known as stress fibers, powered by non-muscle myosin II and crosslinked by α-actinin, are linked to cell-cell and cell-extracellular matrix adhesions and mediate their force-dependent stabilization (8). Branched actin networks become denser and develop the capacity to generate greater forces when exposed to resistive load both *in vitro* (9, 10) and in cells (11), while stress fibers feature mechanical damage repair mechanisms mediated by the force-depedent localization of proteins from the LIN-11, Isl-1 and MEC-3 (LIM) domain superfamily (12, 13). A balance between protrusive forces, generated by branched F-actin, and contractile forces, generated by actin-myosin networks, is necessary for cellular mechanical homeostasis. This is exemplified at adherens junctions, where branched actin pushing facilitates the formation of cadherin-cadherin contacts across adjacent plasma membranes (14, 15), while contractile actin-myosin assemblies are required to maintain intercellular tension and mature junction architecture (16). How distinct network-level force response mechanisms emerge from the activities of individual protein components, many of which overlap between functionally and architecturally distinct networks, remains unclear.

Several ABPs have been reported to engage in force-modulated interactions with individual actin filaments, which could contribute to the force-sensitivity of actin networks (17, 18). This includes canonical F-actin nucleation and polymerization factors (19–24), depolymerization / severing factors (25, 26), and cell adhesion proteins (27–31) whose binding affinity is moderately up- or down-regulated by force, as well as LIM domain proteins (32, 33), which solely bind F-actin in the presence of force. Several force-modulated ABPs feature multiple F-actin binding domains which they use to engage filaments simultaneously, e.g. ARP2/3 (19, 20) and α-actinin (34), suggesting mechanical regulation of their multivalent F-actin binding activities could play a role in the force-sensitive filament crosslinking geometry of networks. Here we specifically focus on the critical cell-cell adhesion protein αE-catenin, which forms both force-stabilized bonds when tension is applied across its F-actin binding interface (“catch bonds”)(27, 30), and preferentially binds mechanically stressed F-actin (“force-activated binding”)(31). The protein exists in two forms, as a soluble dimer with F-actin crosslinking activity that has been suggested to act as a competitive inhibitor of ARP2/3 to coordinate the balance between branched actin generation and actin-myosin contractility at adherens junctions (35), and as a core component of the plasma membrane anchored heterotrimeric E-cadherin–α-catenin–ß-catenin (cadherin-catenin) complex, which serves as the primary linkage between adherens junctions and the cytoskeleton (36). While F-actin catch-bonding by cadherin-catenin complex associated α-catenin has been interpreted to mediate mechanical stabilization of adhesion (37, 38), an activity which is also present in the dimeric form (30), to our knowledge the functional implications of force-activated F-actin binding by the dimeric form have not been assessed. We thus selected dimeric α-catenin to conduct a case study dissecting the interplay between forces, actin network architecture, and multivalent ABP-F-actin binding interactions.

Technical considerations have limited analysis of how force modulates ABP–F-actin binding interactions in the context of higher-order networks. Biophysical assays with single molecule resolution, including optical (23, 25, 27–31) and magnetic tweezers (24) and flow-based systems coupled with total internal reflection fluorescence (TIRF) microscopy (19–22, 26), have provided detailed information into the force sensitivity of ABPs, but these studies have generally been limited to examining individual actin filaments. Analysis of reconstituted actin networks with microrheology techniques (34, 39) and atomic force microscopy (AFM) (9, 10), as well as fluorescence microscopy of mechanically stimulated cells (40), have provided insight into network-level force responses, but generally lack the resolution to report on the behavior of individual ABP–F-actin interactions. Recent studies combining AFM with single-molecule TIRF detected the force-sensitive incorporation of individual ABPs within branched actin networks (9, 10), but the corresponding architecture of the network itself could not be visualized in detail due to nanoscale crowding and the detection limits of light microscopy.

We recently introduced a myosin-motor based force reconstitution assay compatible with TIRF which we have used to characterize force-modulated interactions between ABPs and individual filaments (31, 32). Here, we adapt that assay to the study of networks by incorporating engineered filament end-to-end crosslinking complexes, as well as photo-micropatterning of myosin motor proteins into micron-scale stripes which mimic the sarcomeric organization of stress fibers. We find this system spontaneously generates hundreds of entangled actin networks experiencing myosin-generated forces, in parallel, on a coverslip, facilitating detailed dissection of their architectural dynamics while monitoring ABP engagement. We apply this system to show that myosin forces enhance dimeric α-catenin’s engagement of small filament bundles within networks versus large bundles, likely due to each filament in smaller bundles bearing relatively higher tension than those in larger bundles. Consistently, a mutation which specifically impairs α-catenin’s force-activated actin binding (31) abrogates this effect, resulting in a linear relationship between bundle size and α-catenin binding. We speculate this activity could facilitate localization of α-catenin dimers to contractile networks at newly-formed adherens junctions in order to suppress ARP2/3 activation (35). Our reconstitution system enables systematic dissection of how multivalent ABPs engage architecturally-diverse networks with sufficient throughput to parse inherent heterogeneities; we anticipate it will facilitate exploring mechanistic principles of force-sensitivity that emerge at the network level.

## Results

### Myosins apply tensile force to an engineered paired actin filament complex

Myosin motor proteins are biochemically tractable, which has facilitated the development of fluorescence imaging-based myosin force reconstitution assays for studying mechanically-regulated actin-binding interactions (23, 31–33). In a traditional gliding filament assay (41), a single species of immobilized motor exerts force on filaments, which are translocated across a glass coverslip (Fig. 1A). To capture load-bearing filaments, we previously implemented a “dual motor” system (31, 32), where a mixture of the plus (“barbed”)-end directed motor myosin-5 and the minus (“pointed”)-end directed motor myosin-6 is immobilized, which engage in a tug-of-war. Binding interactions between fluorescently labelled ABPs and F-actin can then be monitored in the presence of ATP-dependent force generation. This system features substantial heterogeneity, as both compressive and tensile forces can be generated depending on the local spatial distribution of motor species along a filament, with force magnitudes that vary with the stochastic progression of motors through their mechanochemical cycles. Thus, although this system has the advantages of utilizing physiological force generators and supporting examination of numerous filaments in parallel, the complexity imposed by its component myosins have limited precise biophysical interpretation of ABPs’ mechanical regulatory mechanisms (31, 32).

**Figure 1.**
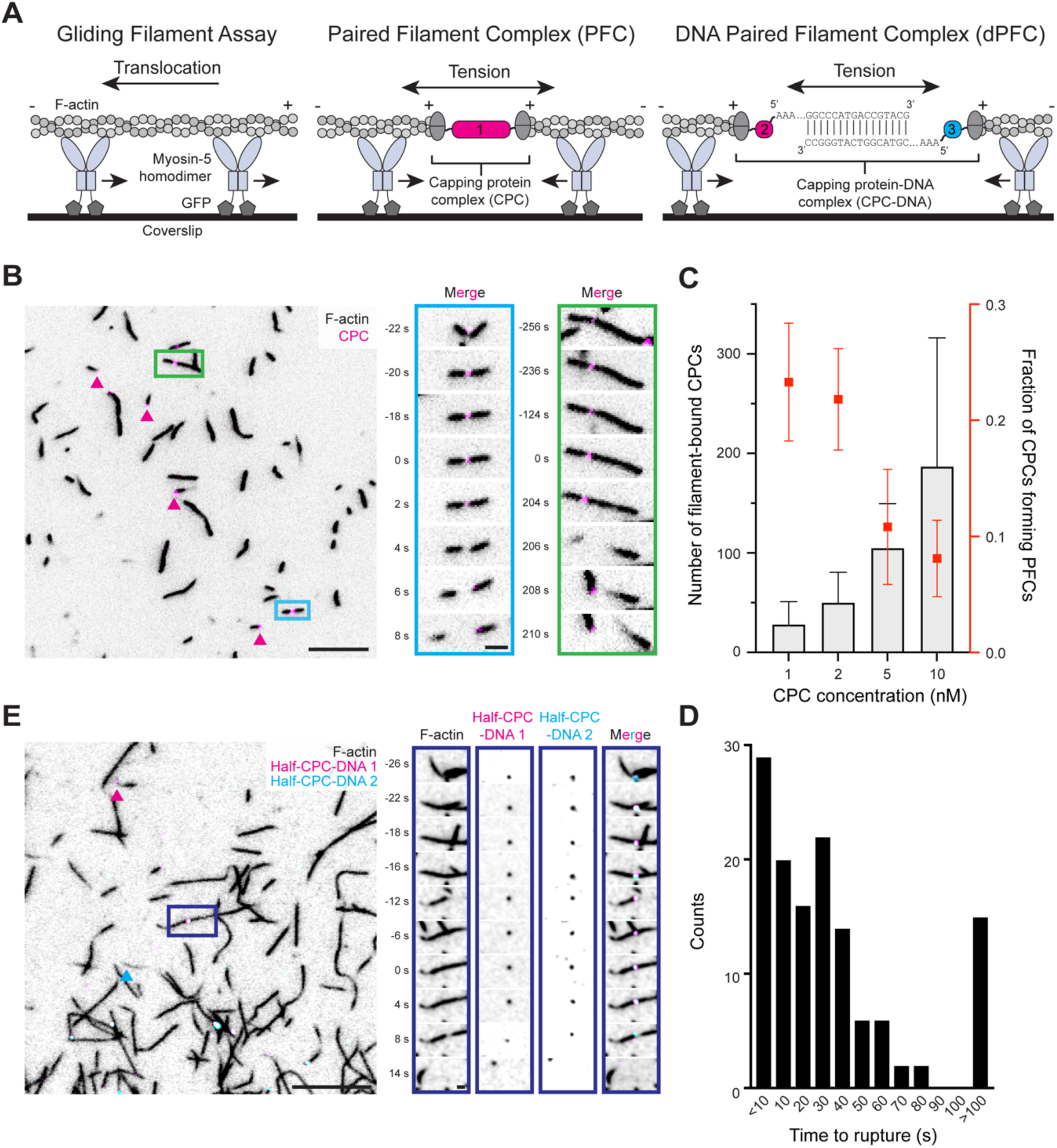
Capping protein complexes assemble actin filaments into paired filament complexes. **A)** Cartoons of a typical gliding filament assay (left), and modified gliding filament assays using CPC to form PFCs (center) or CPC-DNA to form dPFCs (right). **B)** Left: Micrograph of a field of PFCs. Scale bar, 10 μm. Arrowheads indicate single filament complexes. Right: Montages of two PFCs which undergo ruptures. Scale bar, 2 μm. **C)** Quantification of mean number of CPCs bound to filaments (gray bars) versus proportion of CPCs engaged in PFCs (red points) at different CPC concentrations. Error bars represent s.d.; n = 26 from 7 trials. See Fig. S2C for individual measurements. **D)** Quantification of time to rupture of PFCs. n = 130 from 26 trials. **E)** Representative micrograph of a field of dPFCs prepared with CPC-DNA. Scale bar, 10 μm. Arrowheads indicate single filament complexes featuring only one half-CPC-DNA. Right: montage of a PFC where signal from both half-CPC-DNA 1 and 2 are apparent prior to rupture. Scale bar, 2 μm.

With the goal of developing a system featuring a more uniform force distribution, we engineered a complex composed of two copies of the CapZAB capping protein heterodimer, covalently attached through a linker complex. We reasoned this “capping protein complex” (CPC) would engage a pair of filaments by their plus ends, resulting in their head-to-head arrangement (Fig. 1A) in a “paired filament complex” (PFC). In the presence of a plus-end directed motor (myosin-5), PFCs would solely come under tension. The CPC features a modular design which we assemble in a step-wise fashion, using three individually purified protein parts and the Spy/Snoop system (42, 43) to form specific covalent linkages between tagged proteins (Fig. S1A). In our initial assembly, a construct consisting of a HaloTag flanked by two SnoopTags is reacted with an excess of a SpyCatcher-SnoopCatcher adapter. We purify the three-part covalent conjugate by size exclusion chromatography, then react it with an excess of SpyTagged capping protein. A second round of chromatography separates the final five-part complex, which features one HaloTag, two adapters, and two capping protein heterodimers, from intermediates (Fig. S1B). This final purified CPC binds F-actin in a filament end concentration-dependent manner (Fig. S2A).

When mixed with actin filaments labeled with fluorescent phalloidin, CPC generates head-to-head arranged PFCs which come under tension in the presence of immobilized myosin-5 and ATP, which we visualize with TIRF microscopy (Fig. 1B, Movie S1). Formation of PFCs is dependent on the relative abundance of filament ends and CPC. The median filament length in our assay is 8.6 μm. By accounting for typical F-actin structure, we estimate an average of approximately 3,000 actin subunits per filament, and a corresponding barbed end concentration of 0.6 nM. Under conditions where there is a substantial excess of CPC (10 nM), CPCs overwhelmingly associate with single filaments, likely because unbound CPCs have a diffusional advantage in encountering free barbed ends compared with CPCs already associated with a filament on one side. In this regime, we observe numerous CPCs in each field of view, but close to 90% are associated with the trailing end of single gliding filaments as opposed to forming PFCs (Fig. 1C). As CPC concentration is titrated down to 1 nM, there is a steady reduction in the number of singly-filament bound CPCs and a corresponding increase in the proportion of CPCs embedded between two filaments to form a PFC. However, this regime features a lower number of CPCs per field of view (Fig. 1C); thus, there is a trade-off between PFC formation efficiency and the number of productive observations per experiment.

Over time, motors pulling on the two sides of a PFC tear its constituent pair of filaments apart, causing them to explosively separate in opposite directions when the interface between one capping protein heterodimer and its bound actin filament breaks (Fig. 1B, inset). The majority of PFCs persist for dozens of seconds, with a median time to rupture of 26-29 seconds (Fig. 1D) which is independent of CPC concentration (Fig. S2B). A small population of PFCs are long lived, persisting for over 6 minutes before rupturing. While the lifetime of PFC-associated fluorophores exceeds the time to rupture in the vast majority of cases, we do observe instances of the central CPC bleaching in a single step prior to PFC breakage, suggesting we are recording individual CPCs linking filaments (Fig. S2E). Collectively, our engineered CPC generates molecular tugs-of-war across pairs of oppositely oriented single actin filaments which break after tens of seconds under myosin derived tension, sufficient for quantitative observations with TIRF.

### A DNA-based capping protein complex supports modular assemblies

To demonstrate the modularity of our system, and because DNA is a ubiquitous molecular handle with well-defined biophysical properties (44), we next designed and constructed a CPC where the central component is composed of two parts featuring complementary DNA handles that associate through base-pairing (CPC-DNA, Fig. 1A). We assemble this protein-DNA complex using the same SpyTagged capping protein heterodimer, which is conjugated to an adapter featuring a SpyCatcher domain, a HaloTag for fluorescent labelling, and a PCV2 HUH-tag, which forms sequence-specific covalent adducts with DNA (45) (Fig. S1C). This two-part protein complex is then separately reacted with two DNA handles which feature complementary base-pairing regions, via a common specific 5’ DNA sequence which reacts with the PCV2 domain (Fig. S1D). We call the two resultant complexes half-CPC-DNA 1 and half-CPC-DNA 2. Sequential rounds of size exclusion chromatography purify the protein-DNA adducts, followed by anion exchange chromatography to remove residual protein which is not DNA conjugated.

Half-CPC-DNA 1 and half-CPC-DNA 2 are each able to independently engage and translocate with the plus ends of actin filaments in gliding filament assays (Movie S2, S3). Mixed overnight, the two half-CPC-DNAs become base paired, forming a full CPC-DNA. When CPC-DNA is added to F-actin, it is capable of assembling filaments into DNA-based PFCs (dPFCs) which break under tensile force (Fig. 1E and S3A, Movie S4), as we observed with PFCs assembled with the proteinaceous CPC (Fig. 1B). We observe that, upon rupture, both fluorescently-labelled half-CPC-DNAs remain associated with the same filament, indicating the breakage occurs at the interface between a capping protein heterodimer and its bound filament. Notably, dPFC assembly occurs at lower efficiency with CPC-DNA compared to PFC assembly with the proteinaceous CPC, and we see numerous instances of single half-CPC-DNAs (either half-CPC-DNA 1 or half-CPC-DNA 2) translocating in isolation associated with filament plus ends, as well as full CPC-DNAs associated with single filaments (Fig. S3A and B), suggesting this system will require further optimization for practical experimental utility. Nevertheless, successful dPFC assembly with CPC-DNA demonstrates the generalizability of our CPC approach: such an arrangement may prove useful for the integration of DNA based fluorescent force-reporters (46) or to generate defined higher-order assemblies of tensed filaments through DNA origami and sequence-addressable handles (47).

### Micropatterning myosin motors generates higher-order PFC networks

While individual PFCs displayed the anticipated behavior in the presence of barbed-end directed force, the infrequency of spatially isolated PFCs on stochastically distributed fields of myosin-5 limited their utility for investigating mechanically-regulated ABP-F-actin interactions. Inspired by the sarcomeric organization of contractile muscle fibers (48) and stress fibers (8), we implemented a protein micropatterning approach, using a PRIMO device to photo-pattern 10 μm-spaced stripes of fibrinogen-GFP binding protein (49) that is specifically bound by GFP-tagged myosin-5, interleaved with 10 μm passivated gaps (Fig. S4). In this geometry, we reasoned PFCs would become trapped between stripes as motors on adjacent stripes pull paired filaments in opposite directions, while unpaired single filaments would be attracted onto the stripes. Despite this spatial organization, the system still features heterogeneities. Filaments can be engaged by variable and fluctuating numbers of motors, whose positions remain random within the stripes, and thus we anticipate force magnitude will vary both between different filaments and along individual filaments in time. This configuration nevertheless recapitulates some of the mechanical complexity of cytoskeletal networks in cells, as the underlying forces are generated by physiological motors, whose variability in quantity, synchronization and inter-motor spacing also impose a range of forces on F-actin *in vivo*. On the addition of F-actin pre-incubated with CPC and ATP to our micropatterned surface, the vast majority of F-actin binds to the motor protein stripes. Over time, single filaments protruding from the edge of stripes are either pulled onto them or pushed off and released into solution as the motors operate (Fig. 2A). A sparse collection of filaments remain spanning the gaps between stripes, which we interpret to be filaments in PFCs which become trapped as motors on adjacent stripes pull the two sides of the PFC in opposite directions. Consistently, in the absence of CPC, the limited number of filaments which remain spanning the gaps between stripes do not move, suggesting they are trapped by binding inactive motors or adhering to the coverslip on one side (Fig S5A), while samples featuring PFCs continue to undergo slow force-dependent dynamics. This self-organization is in marked contrast to the unordered dynamics of PFCs and single filaments on stochastically distributed fields of motors (Fig. 2A).

**Figure 2.**
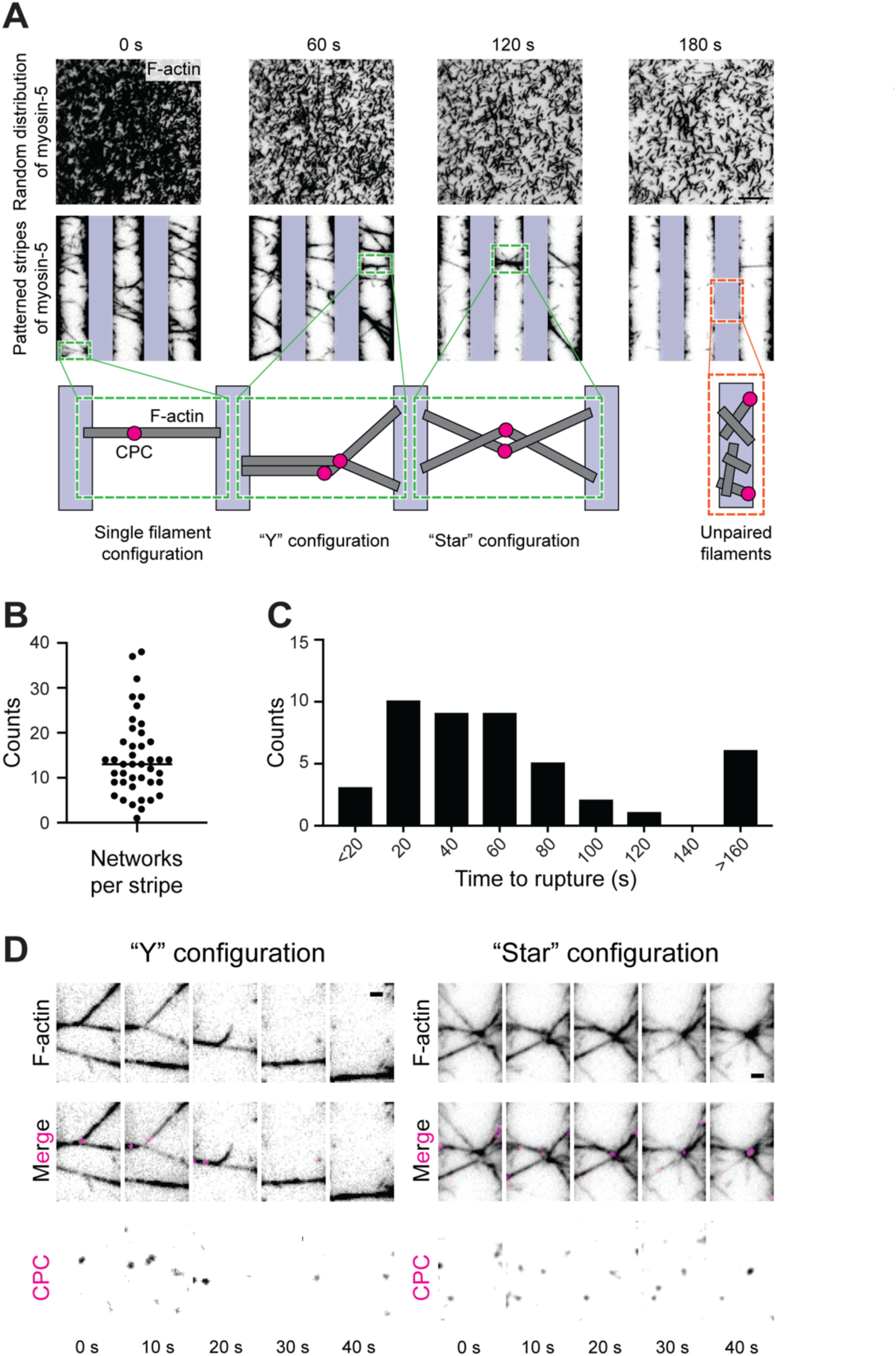
Micropatterning myosin-5 collects PFCs into tensed networks. **A)** Montages of PFCs on stochastically distributed (top) and micropatterned (bottom) fields of myosin-5. F-actin is black; CPC is not shown. Vertical bars indicate positions of myosin-5 stripes. Scale bar, 10 μm. Inset cartoons indicate PFC configurations at highlighted positions. **B)** Quantification of number of PFC networks per stripe. Bar indicates mean. n = 42 from 4 trials. **C)** Quantification of times to rupture of PFC networks. n = 45 from 4 trials. **D)** Montages of “Y” configuration (left) and “star” configuration (right) networks breaking over time. Scale bars, 2 μm.

Unlike the single PFCs observed on stochastic motor fields (Fig. 1B), we primarily observe collections of PFCs becoming tangled as the motors operate, generating interwoven networks of tensed bundles. These complex network structures are highly dynamic, rearranging as their constituent filaments break before motor forces ultimately result in their complete destruction (Movie S5). The flexible head-to-head filament orientation enforced by the CPC combined with motor micropatterning is necessary for networks to emerge; when either of these aspects of the system is absent, networks do not form. In the absence of CPC, the few filaments that are trapped between stripes do not interweave to form higher order networks, persisting instead as individual filaments / small bundles with lower F-actin fluorescence intensities than CPC-dependent networks (Fig. S5A-C). As no additional filament crosslinking ABPs are included in our assay, the collection of filaments into bundles emerges solely from tensile forces acting on webs of filaments entangled by the CPC. We observe dozens of networks in each microscopic field of view (Fig 2B), which vary in size, geometric complexity, and lifetime, with most networks persisting for tens of seconds to minutes (Fig. 2C).

The filament arrangement enforced by the CPC leads us to infer that bundled filaments within networks are predominantly parallel, with filament plus ends congregating around the central knot. We note two prominent arrangements of such knots, which we refer to as the “Y” and “star” configurations (Fig. 2A). In the Y configuration, at least two PFCs are entwined to generate a three-pronged structure. This architecture facilitates observing the dynamic force-balance imposed by the motor stripes while networks rearrange, as breakage of one prong results in the joint between the other two straightening, thereby producing a colinear bridge between stripes (Fig. 2D). In “star” configurations, filaments radiate outwards from a central knot, where CPCs tend to be gathered tensile forces over time (Fig. 2E). We infer this to be the result of the greater deformation tolerance of CPCs, which feature multiple flexible linkers within their covalent assembly, versus actin filaments, whose non-covalent inter-subunit interfaces can be readily broken by motor forces under our assay conditions (32). Star configurations are prone to sequential rearrangements as their constituent prongs break, drastically changing the shape of the structure over time before it collapses. Notably, in both network configurations, the prongs composing networks have widely varying fluorescence intensities, indicating that they are primarily composed of bundles rather than individual filaments. In sum, combining PFCs with motor micropatterning enables us to generate dozens of tensed networks in parallel with varying dynamic architectures, providing a platform to probe the interplay of tension and network architecture in controlling F-actin engagement by ABPs.

### Dimeric α-catenin features force-activated F-actin binding

We next employed our system to dissect how force and network architecture impact F-actin engagement by dimeric α-catenin. The protein’s dimerization occurs via a domain in the protein’s N-terminal head, while the C-terminal tail contains its actin-binding domain (ABD). Here we employ a construct spanning amino acids 56-906 of human α-catenin with an N-terminal HaloTag for fluorescent labelling, where the first 55 N-terminal residues were truncated to enhance stability. The purified protein eluted from a size exclusion chromatography column at a retention volume consistent with dimerization (Fig. S6A). As this construct comprises the majority of the α-catenin sequence, we hereafter refer to it as “full-length” (FL) α-catenin for simplicity. As our previous studies of α-catenin’s mechanosensitivity focused on its isolated ABD, we first examined whether dimeric FL α-catenin maintains force-activated F-actin binding activity. We previously found the ABD’s force-activated binding depends on the protein’s disordered C-terminal extension (CTE) comprising residues 872-906, as removal of this segment compromised increased binding to tensed F-actin (31). We thus generated an analogous construct in FL α-catenin, comprising residues 56-871, which we refer to as α-cateninΔC. This construct also eluted as a dimer from a size exclusion column, and both FL α-catenin and α-cateninΔC displayed F-actin binding and bundling activity in co-sedimentation assays (Fig. S6B).

We next examined the force-dependent F-actin engagement of both constructs in our assay by comparing their F-actin association before and after adding ATP to activate motors, conducting paired analysis of the same microscopic fields. Upon addition of ATP, qualitative changes in filament orientation and morphology occurred, as slackened filaments are pulled taut by the motors and interdigitated networks undergo force-dependent rearrangements (Fig. S7A, Movie S6). Ratiometric measurement of the fluorescence intensity of bound FL α-catenin versus underlying F-actin revealed a marked increase in the per-filament binding of FL α-catenin in the presence of ATP, a behavior which was essentially uniform across trials, demonstrating this form of α-catenin also possesses force-activated binding activity. Notably, an increase in the α-catenin : F-actin ratio indicates enhanced binding on the per-filament level, rather than the signal increase being driven by filaments being incorporated into larger bundles through motor activity (Fig. S7B). Consistently, we do not observe a significant change in the distribution of F-actin intensities upon ATP addition (Fig. S7C). Conversely, α-cateninΔC displays a more variable response, without a striking increase in F-actin binding upon ATP addition (Fig. S7B). We do nevertheless observe a significant increase (p = 0.011) in its binding on average, suggesting the CTE deletion impairs but does not completely eliminate force-activated binding. Notably, we were able to observe this weak effect due to the substantially higher throughput enabled by the PFC / motor patterning assay presented here versus our prior implementation of the dual-motor assay (31). Nevertheless, our data are broadly consistent with dimeric FL α-catenin possessing force-activated actin binding activity mediated by the protein’s CTE, suggesting this occurs through a similar mechanism as we previously reported for the isolated α-catenin ABD (31).

### The distribution of α-catenin is heterogeneous within individual networks

To assess how α-catenin’s force-activated binding could intersect with its engagement of higher-order F-actin assemblies, we undertook a detailed analysis of a large network composed of several interlocking stars (Fig. 3) in the presence of ATP and FL α-catenin. Qualitatively, we observed substantial heterogeneity in both the F-actin and α-catenin fluorescent intensities across the different prong-like segments composing the network (Movie S7). As the network underwent mechanical transitions prior to its eventual collapse, we noted substantial changes in α-catenin intensity in specific segments. To quantify this phenomenon, we time averaged frames between substantive transitions (when constituent filament bundles break or change position), which we refer to as “states”, manually split the network into its segments, and measured the average fluorescence intensity of both α-catenin and F-actin in each segment across states (Fig. 3A).

**Figure 3.**
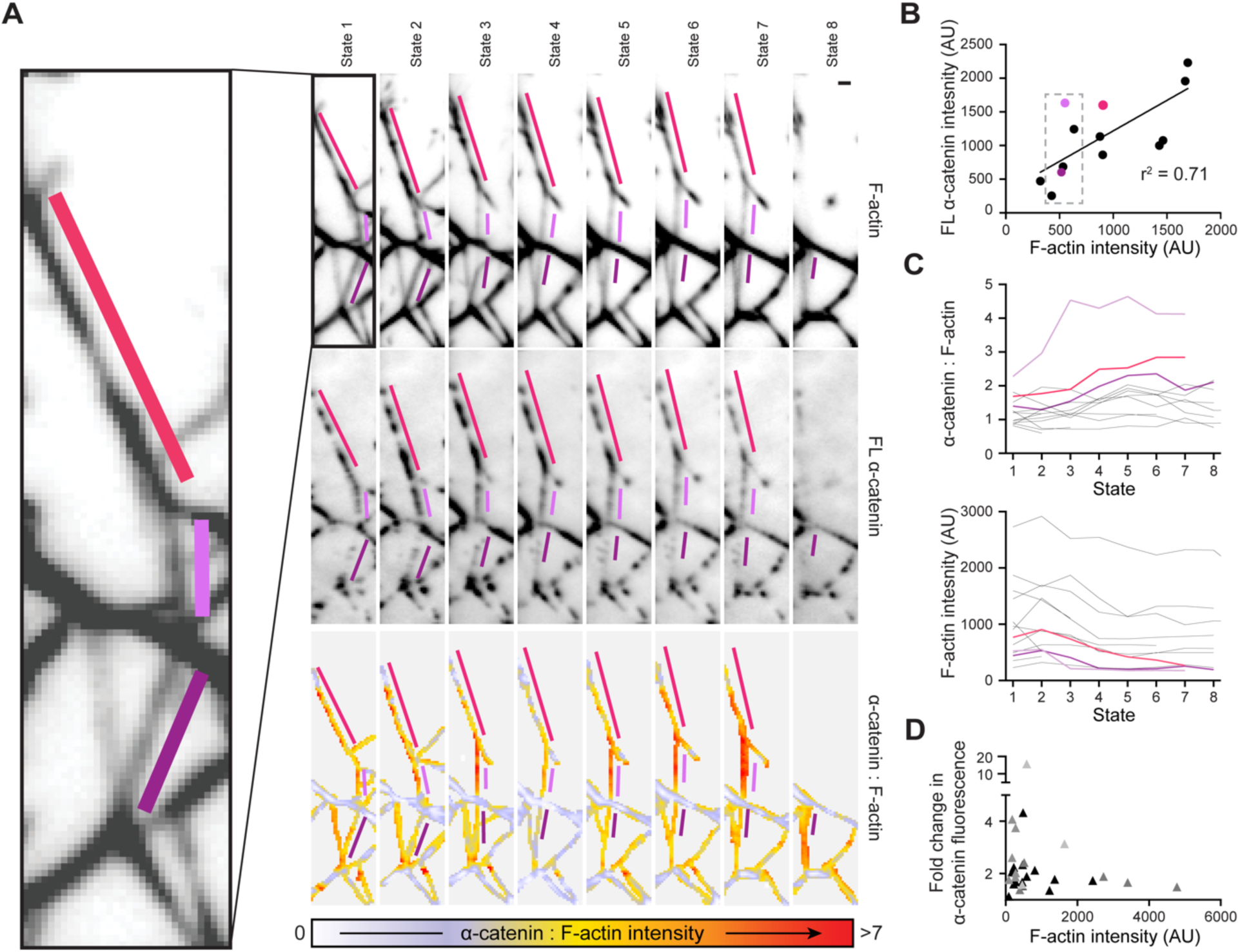
FL α-catenin binding is heterogenous within tensed F-actin networks. **A)** Time averaged images of actin fluorescence intensity (top), FL α-catenin fluorescence intensity (middle) and α-catenin : F-actin fluorescence intensity ratio (bottom) of 8 configurationally distinct states of a PFC network. Bundle segments highlighted in shades of pink feature an increase in their α-catenin : F-actin fluorescence intensity ratio over time. Scale bar, 2 μm. **B)** Scatterplot of F-actin versus α-catenin fluorescence intensity of bundle segments from state 2 in **A**. Black points represent non-highlighted bundle segments. Linear regression is displayed. **C)** α-catenin : F-actin fluorescence intensity ratio (top) and F-actin fluorescence intensity (bottom) of bundle segments across all 8 states of the network. Black lines represent non-highlighted bundles segments. **D)** Scatterplot of maximal fold-change in α-catenin fluorescence intensity during the imaging period versus F-actin intensity of bundle segments. n = 31; shades of grey indicate measurements from the 4 networks analyzed.

We first examined the relationship between segment α-catenin intensity and F-actin intensity, focusing on state 2 (Fig. 3B). While there was a moderate correlation between the amount of F-actin in a segment and the degree of α-catenin binding, segments with similar F-actin intensity could feature α-catenin binding which varied by greater than 5-fold (Fig. 3B). This suggests that in the context of mechanically active networks, FL α-catenin’s binding is not solely controlled by the local density of F-actin binding sites. When monitored across states, multiple segments featured a dynamic increase in their α-catenin : F-actin ratio (Fig. 3C). This was not due to an increase in F-actin abundance, as there is no excess G-actin in our system to support additional polymerization, and all segments featured a loss of F-actin signal over time (Fig. 3C), likely due to filament breakage and fluorophore bleaching. The most parsimonious explanation for this observation is that a subset of segments within the network are more susceptible to force-enhanced α-catenin binding, which reweights the distribution of α-catenin across the network as it undergoes mechanical rearrangements.

### α-catenin’s force-activated F-actin binding is more prominent in smaller actin bundles

We next sought to determine the mechanistic basis of α-catenin’s heterogeneous force-activated binding within networks. We noted that in the representative network we examined, segments which featured a dynamic increase in α-catenin binding across states tended to have lower F-actin intensities relative to segments featuring stable α-catenin binding (Fig. 3C), leading us to hypothesize that smaller bundles could be more susceptible to mechanical regulation. Consistently, when examining data from three additional networks, we observed that smaller network segments qualitatively appeared to feature a broader distribution in the fold change of their α-catenin intensities over time (Fig. 3D). To quantitatively examine whether this phenomenon was mediated by α-catenin’s force-activated actin binding activity, we compared the relationship between network segment size (F-actin intensity) and α-catenin engagement (α-catenin intensity) for both FL α-catenin and α-cateninΔC in the presence and absence of myosin force generation. To facilitate higher-throughput analysis, we conducted unpaired comparisons of specimens treated with ATP versus those treated with apyrase to remove residual nucleotide from the system. Networks were automatically detected and segmented by binarization and thresholding of the F-actin channel (Fig. S8B), allowing us to examine thousands of segments across experimental replicates (Fig. 4A).

**Figure 4.**
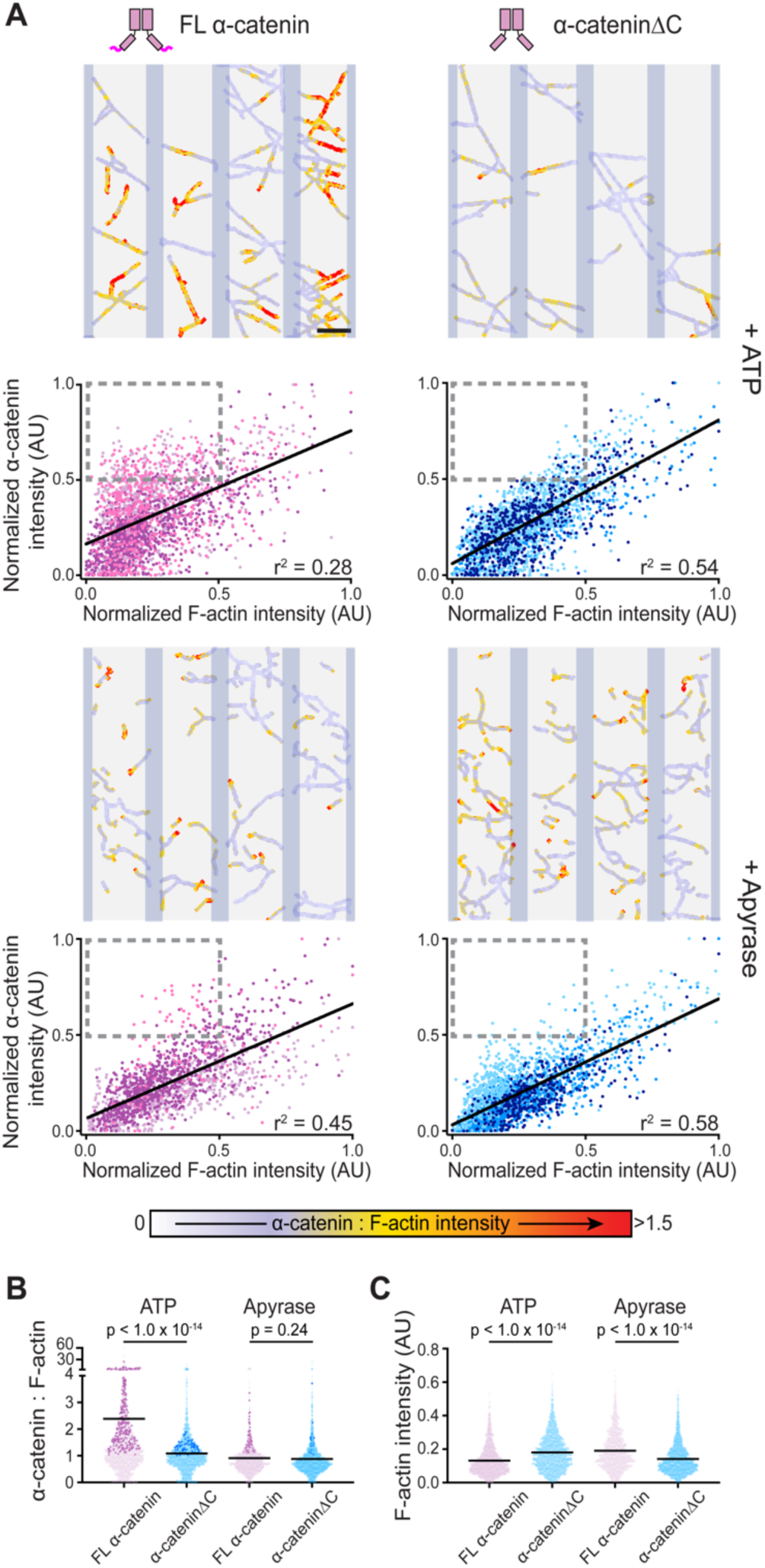
Force-activated F-actin binding mediates FL α-catenin’s enrichment on small bundles. **A)** Representative α-catenin : F-actin fluorescence intensity ratio images of PFC networks (vertical bars indicate positions of myosin-5 stripes) and associated scatter plots of normalized F-actin versus α-catenin fluorescence intensity of bundle segments for FL α-catenin (left) or α-cateninΔC (right), in the presence of ATP (top) or apyrase (bottom). Different colored points indicate data from 3 independent trials. n = 2,449 (FL α-catenin + ATP); 2,243 (FL α-catenin + apyrase); 1,668 (α-cateninΔC + ATP); 2,241 (α-cateninΔC + apyrase). Grey boxes indicate quadrant representing the top 50% of α-catenin fluorescence intensity and bottom 50% of F-actin fluorescence intensity. Linear regressions are displayed. Scale bar, 10 μm. **B)** Pooled analysis of α-catenin : F-actin fluorescence intensity ratios of data from **A** compared with unpaired Welch’s t-test. Bars indicate means; darker points correspond to boxed quadrants in **A**. **C)** Pooled analysis of F-actin fluorescence intensity of data from **A**, analyzed as in **B**.

In the apyrase (no force) condition, both FL α-catenin and α-cateninΔC displayed an approximately linear relationship between F-actin intensity and α-catenin intensity, suggesting in this regime network engagement is driven purely by mass action without dimeric α-catenin’s multiple actin-binding domains conferring notable avidity effects. However, in the presence of ATP and motor activity, FL α-catenin’s distribution broadens, with a notable enrichment of segments in the quadrant representing the upper 50% of α-catenin intensity and the lower 50% of F-actin intensity (Fig. 4A, grey dotted rectangle). This effect is absent in the α-cateninΔC force-sensing deficient mutant, suggesting it is specifically mediated by α-catenin’s force-activated F-actin binding activity. This subpopulation of smaller bundles extensively bound by α-catenin contributes markedly to FL α-catenin’s significantly greater mean α-catenin : F-actin ratio in the +ATP condition (Fig. 4B, darker points), driving the phenomenon of force-activated binding at the bulk level. While we do observe significant differences in the distribution of F-actin intensities between conditions, these are modest and do not clearly track with force-activated binding by FL catenin in the +ATP condition (Fig. 4C). Collectively, these data suggest that in the presence of myosin activity dimeric FL α-catenin’s force-activated binding enhances binding to smaller bundles, leading to an equalization of total α-catenin across networks of different sizes.

## Discussion

Here, we introduce a reconstitution platform that allows the dissection of force-regulated ABP binding behaviors in the context of higher-order F-actin networks. Our approach enables the heterogeneous architecture and dynamics of micron-scale networks experiencing active myosin forces, recapitulating aspects of the complexity of actin networks in cells, to be analyzed in quantitative detail with TIRF microscopy. While our proteinaceous CPC / PFC system spontaneously forms heterogenous networks on micropatterned substrates, the CPC-DNA system can likely be further engineered to facilitate the generation of defined assemblies through DNA nanotechnology (47). Additionally, DNA-based fluorescent force-reporters (46) should be compatible with this system, which would enable examining the relationship between local force magnitudes and ABP engagement, of particular importance for establishing detailed mechanisms of force-regulated binding. Our myosin micropatterning approach complements previously reported reconstitution systems where actin nucleation factors are patterned in the presence of soluble motors (50, 51), which also give rise to active networks with emergent micron-scale dynamics. More complex patterns of motors, as well as simultaneous patterning of motors and actin nucleating factors, may facilitate reconstitutions of increasing complexity that recapitulate multiple types of actin networks simultaneously side-by-side, enabling studies of how ABPs are partitioned across networks within a common compartment.

We applied our approach to study the force-regulated binding of dimeric FL α-catenin, finding that, in the presence of myosin forces, the protein preferentially engages smaller actin bundles embedded within higher order networks. Our data are compatible with a model in which these network segments bear greater load per filament than larger bundle segments (Fig. 5), rendering them more susceptible to force-evoked conformational transitions which modulate α-catenin engagement. This mechanism presupposes that motors apply force of a similar total magnitude to a network segment regardless of its size, a criterion which is likely fulfilled by the embedded segments analyzed in Fig. 3, which do not directly contact the motor stripes and thus experience forces distributed through the network. A non-exclusive alternative is that the filament packing of larger bundles could potentially also restrict the conformational landscapes of their component filaments. A limitation of our study which precludes discriminating between these mechanisms is our inability to directly measure the distribution of forces, which can likely be overcome in future work using force probes embedded in CPC-DNA as described above. Our system is furthermore in principle compatible with cryo-electron microscopy studies, which could be used to examine the conformational landscape of F-actin within tensed PFC bundles.

**Figure 5.**
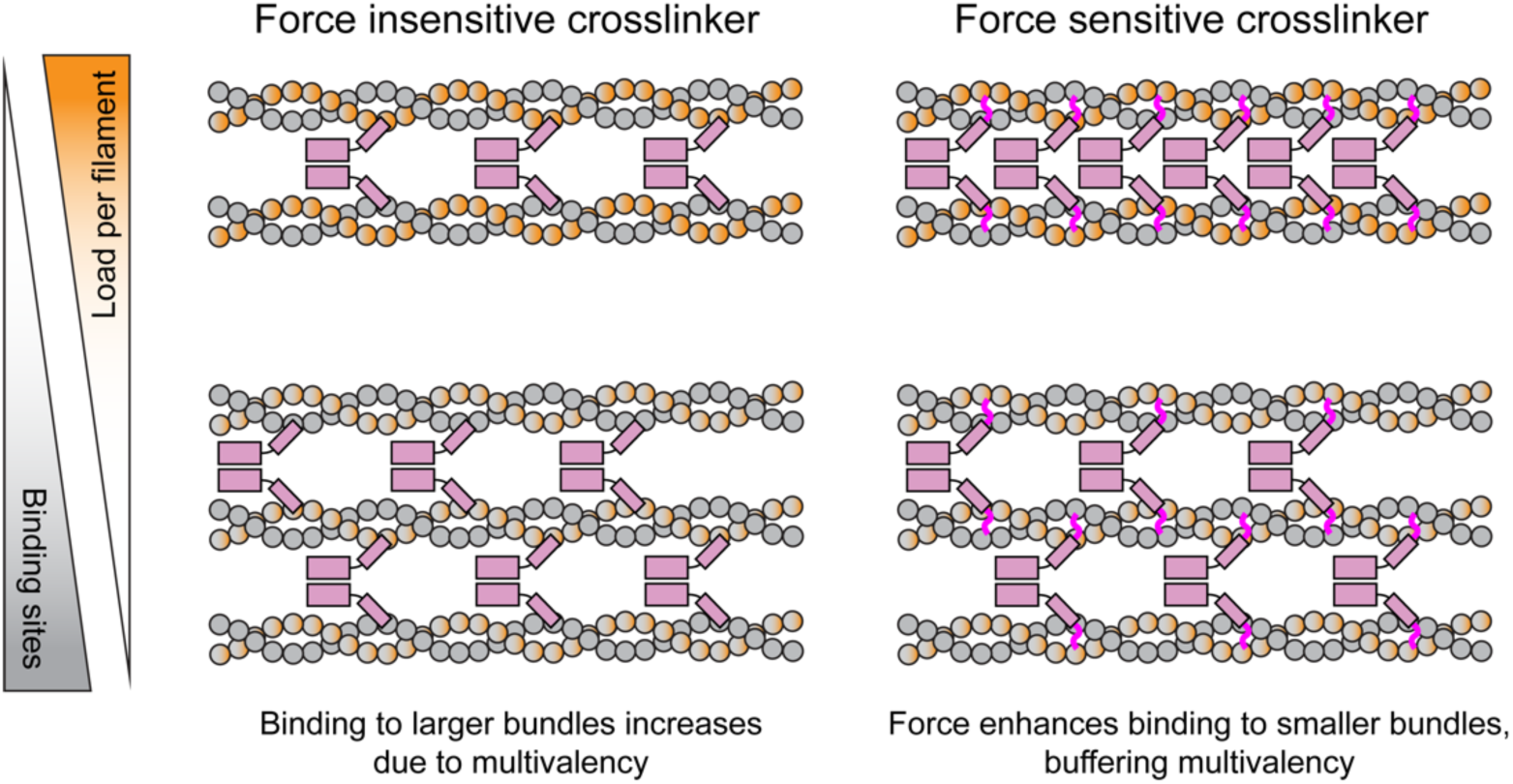
Conceptual model of force-mediated bundle size discrimination. F-actin crosslinkers with force-activated F-actin binding activity balance enhanced engagement of smaller filament bundles, where individual filaments experience a higher load per filament, with mass-action driven association to larger bundles featuring more binding sites. This balances the total number of crosslinkers per bundle across a range of filament bundle sizes. Bundle engagement by force insensitive F-actin crosslinkers is driven solely by the number of binding sites, leading to their accumulation on larger filament bundles.

The absence of clear avidity effects conferred by dimeric α-catenin’s dual actin-binding domains is unexpected, as actin-bundling proteins are anticipated to display enhanced engagement of multi-filament assemblies. A recent structural study of FL α-catenin suggested that both of the protein’s ABDs can simultaneously bind the same actin filament (52), mediated by the well-established flexibility of the head domain and its linkage to the ABD (53). As we nevertheless clearly observe actin-bundling activity (Fig. S6B), as has previously be reported (54), our data are compatible with a model in which the protein is capable of binding single filaments with both ABDs as well as bridging pairs of filaments. In this scenario, the protein’s binding should be driven solely by the total availability of actin binding sites rather than their arrangement into a specific geometry, which would increase linearly with bundle size. As only the ABD of FL α-catenin bound to F-actin was visualized by Izard and colleagues (52), future structural studies will be required to dissect the protein’s potential multiple F-actin binding modes.

Biologically, we speculate that α-catenin’s force-activated binding of smaller F-actin bundles could facilitate equalizing the protein’s distribution across bundles of differing sizes in cells. This could potentially mediate inhibition of ARP2/3 branching by dimeric α-catenin as contractile networks are formed at nascent adherens junctions (35), a hypothesis which could be tested in future work by examining the junctional dynamics of epithelial cells expressing the force-activated binding deficient mutant α-cateninΔC. Other force-activated actin binding proteins have been reported to display mechanically triggered localization to smaller F-actin bundles embedded within larger networks in cells. Zyxin (12), along with other LIM proteins (13, 32, 33), rapidly localizes to “stress fiber strain sites”, mechanically-induced tears in stress fibers which feature reduced actin density (12), suggesting that the mechanical fragility of network segments featuring low filament numbers may be more generally productively harnessed by cells. The tools we introduce here should more broadly facilitate investigating how this and other network-level phenomena intersect with mechanical regulation of ABPs.

## Acknowledgements

We gratefully acknowledge Pinar Gurel and Harry Higgs for the gift of the CapZA1 / CapZB constructs. We also acknowledge Wendy Gordon for the gift of the PCV2 construct and fruitful discussions of its use. We also acknowledge Scott Gradia for the gift of the 2HRT pET vector (via Addgene, plasmid # 29718). This work was funded by a Pew Biomedical Scholar Award, an Alfred P. Sloan Foundation Matter-to-Life Award (G-2020-14047), and NIH / NIGMS grant R01GM141044 to G.M.A.

## Author Contributions

JL and GMA designed research. JL, AP, and MB performed research. JL, AP, and GMA analyzed data. JL and GMA wrote the paper, with input from all authors.

## Data and Materials Availability

TIRF movies and raw data supporting this study will be made available at DOI: 10.5281/zenodo.8239706. All DNA constructs are available from the corresponding author without restriction.

## Code Availability

Custom scripts used to quantify TIRF data will be made available at: https://github.com/alushinlab/patterns-PFCs

## Methods

### Actin purification

Actin was purified from chicken skeletal muscle as described previously (32, 55), and all steps were conducted at 4°C. Briefly, chicken muscle acetone powder was resuspended in 20 mL of G-Ca buffer (2 mM Tris-Cl pH 8.0, 0.5 mM DTT, 0.2 M ATP, 0.01% NaN_3_, 0.1 mM CaCl_2_) and mixed by inversion for 30 min before centrifugation at 185,913 g in a Ti70 rotor for 30 min. The supernatant was filtered through 90 mm filter paper (Whatman) and retained. The pellet was then resuspended in another 20 mL of G-Ca buffer and the inversion mixing, centrifugation, and filtering steps were repeated. Next, 50 mM KCl and 2 mM MgCl_2_ were added to the pooled supernatants, which contain actin monomers, to initiate actin polymerization, followed by incubation for 1 hour. Dry KCl was then added to achieve a concentration of 800 mM and the solution was incubated for 30 min to facilitate the dissociation of contaminating factors from actin filaments. The solution was then divided into two tubes and centrifuged in a Ti70 rotor at 185,913 g for 3 hours. Each pellet was resuspended in 2 mL of G-Ca buffer and incubated overnight.

The resuspended pellets were gently transferred to a Dounce chamber, homogenized for 30 passes, sheared three times through 26G and 30G needles consecutively, then dialyzed in SnakeSkin dialysis tubing (MWCO 10 kDa) (Thermo Fisher) in 1 L of G-Ca buffer overnight. The actin solution was then sheared through a 30G needle again before dialysis in 1 L of fresh G-Ca buffer for another day. It was then centrifuged in a Ti90 rotor at 419,832 g for 3 hours. The upper 2/3 of the supernatant was purified by size-exclusion chromatography using a HiLoad 16/600 Superdex 200 column (Cytiva). The second half of the peak was collected and purified actin was maintained in G-Ca buffer at 4°C before use.

### F-actin preparation

F-actin was polymerized fresh for each experiment from G-actin in G-Mg (2 mM Tris-Cl pH 8.0, 0.5 mM DTT, 0.2 M ATP, 0.1 mM MgCl_2_, 0.01% NaN_3_) buffer with KMEI (50 mM KCl, 1 mM MgCl_2_, 1 mM EGTA, 10 mM imidazole pH 7.0, 1 mM DTT) added to initiate polymerization as described previously (56). Polymerization was left to occur at room temperature for 1-2 hours. For TIRF microscopy, polymerized F– actin was incubated with fluorescently labelled phalloidin (Alexa Fluor™ Plus 555 Phalloidin or Alexa Fluor™ 488 Phalloidin, Invitrogen) in a 1:1.2 (actin:phalloidin) molar ratio for 10 min at room temperature before being placed on ice.

### Expression cloning

Expression vectors for α-catenin were constructed by inserting the cDNA sequence of *H. sapiens* αE-catenin_56-906_ (FL α-catenin) or αE-catenin_56-871_ (α-catenin ΔC) and HaloTag (Promega) into pET vector 2HR-T through Gibson assembly (57).

Expression vectors for CPC components were constructed by insertion of synthesized *E. coli* codon optimized cDNA sequences into bacterial expression vectors with N-terminal His6-tags and 3C cleavage sites by Gibson assembly. pET vector 2HR-T was used for the SpyCatcher-SnoopCatcher, SnoopTag-HaloTag-SnoopTag, SpyCatcher-HaloTag-PCV2 constructs. A pRSF Duet 1 vector (Millipore Sigma) was used for the CapZA and SpyTag-CapZB construct. CapZA and CapZB cDNA sequences were derived from *M. musculus* CapZA1 (286 residues) and CapZB (isoform 2, 272 residues). cDNA sequences for SpyTag, SpyCatcher, SnoopTag amd SnoopCatcher were identical to those reported by Howarth and colleagues (42, 43).

Expression vectors for myosin-5 and calmodulin were previously described (32). Briefly, for myosin-5, *H. sapiens* myosin 5a (residues 1–1091) was cloned into a modified pCAG mammalian expression vector with C-terminal non-fluorescent GFP-tag (S65T) and a Flag-tag. For calmodulin, human full-length calmodulin (residues 1-149) was cloned into a modified pCAG mammalian expression vector with no tag.

### Expression and purification of proteins

α-catenins and CPC components were expressed in BL21(DE3) *E. coli* cells (New England Biolabs), grown in LB media at 37°C to an optical density of 0.5-0.8 and induced with 0.2 mM IPTG. After induction, the cells were grown overnight at 15°C, spun down at 15,970 g in a JLA 8.1 rotor (Beckman Coulter), and cell pellets collected and stored at −80°C until use.

Cell pellets were resuspended in lysis buffer (50 mM Tris-Cl pH 8.0, 150 mM NaCl, 2 mM β-mercaptoethanol, 20 mM imidazole) containing lysozyme (Affymetrix/USB) at a 1 mg/mL working concentration, nutated at 4°C for 1 hour, after which the lysate was sonicated for 5 min. The lysate was then clarified by spinning in a JA 25.50 rotor (Beckman Coulter) at 48,380 g for 30 min at 4°C. Cleared lysate was incubated with Ni-NTA resin (Qiagen) for 2 hours on a rotator at 4°C, after which the flow-through was discarded and the resin was washed with two bed volumes of wash buffer (50 mM PO_4_^3−^ pH 8.0, 150 mM NaCl, 20 mM imidazole, 2 mM β-mercaptoethanol). Proteins were subsequently eluted in elution buffer (50 mM PO_4_^3−^ pH 8.0, 150 mM NaCl, 250 mM imidazole, 2 mM β-mercaptoethanol). For α-catenins, purified His-tagged TEV protease (prepared according to a published protocol (58)) was added at 0.05 mg/mL working concentration, then the eluent was dialyzed against dilution buffer (50 mM PO_4_^3−^ pH 8.0, 150 mM NaCl, 2 mM β-mercaptoethanol) overnight, during which time digestion occurred. For CPC components, purified His-tagged 3C protease (prepared according to a published protocol (59)) was added at 0.05 mg/mL working concentration, then similarly dialyzed. These mixtures were incubated with Ni-NTA resin and the flow-through containing cleaved protein was retained. Ion exchange chromatography was conducted on a HiTrapQ anion exchange column (Cytiva) followed by size exclusion chromatography on a Superdex 200 Increase column (Cytiva) in gel filtration buffer (50 mM PO_4_^3−^ pH 8.0, 150 mM NaCl, 2 mM β-mercaptoethanol, 1 mM EDTA), and the final protein concentration was estimated using the Bradford colorimetric assay (Pierce), calibrated with BSA (Gemini). Lastly, 10% v/v glycerol was added, and the protein snap-frozen in liquid nitrogen and stored at −80°C until use.

Myosin-5 was purified from transiently transfected FreeStyle HEK293 cells (Thermo Fisher) as previously described (32). Briefly, HEK cells were grown to a density of 1×10^6^ cells/ml before co-transfection with myosin-5 and calmodulin. Calmodulin and myosin-5 expression plasmids were prepared from *E.coli* 5α cells (New England Biolabs), transformed with the each of plasmids, grown up overnight in a 500 ml culture and maxiprepped (PureYield Maxiprep Kit, Promega). 400 μg of calmodulin and myosin-5 expression vectors (at a 1:6 ratio) were mixed with 15 ml FreeStyle 293 media (Thermo Fisher) and 1.2 ml of PEI (Polysciences) before incubation for 15 min at room temperature, followed by transfection by mixing with 400 mL of cells. Transfected cells were grown on an orbital shaker at 250 rpm in 8% CO_2_ at 37°C, then harvested 72 hours post transfection by centrifugation at 15,970 g in a JLA 8.1 rotor (Beckman Coulter). Cell pellets were snap-frozen in liquid nitrogen and stored at −80°C until use.

For purification, cell pellets were resuspended in lysis buffer (50 mM Tris-HCl pH 8.0, 150 mM NaCl, 2 mM MgCl_2_, 0.2% CHAPS, 2mM ATP, 1 mM PMSF, 1 mg/mL aprotinin, leupeptin, and pepstatin), and were incubated on a rocker for 1 hour. The lysate was clarified by spinning in a JA 25.50 rotor (Beckman Coulter) at 48,380 g for 30 min at 4°C. The resultant supernatant was mixed with anti-Flag M2 affinity beads (Sigma-Aldrich) and incubated on a rocker for 1.5 hours. The protein-bound beads were washed three times with wash buffer (50 mM Tris-HCl pH 8.0, 150 mM NaCl, 2 mM MgCl_2_, and 2 mM ATP). Protein was eluted with wash buffer supplemented with 100 mg/mL Flag peptide (Sigma-Aldrich). The eluent was buffer-exchanged to a storage buffer (10 mM Tris-HCl pH 8.0, 100 mM NaCl, 2 mM MgCl_2_, and 3 mM DTT) using a spin concentrator (Amicon Ultra-4, MWCO 50 kDa), snap-frozen in liquid nitrogen and stored at −80°C until use.

Human calmodulin was also purified from Bl21(DE3) *E. coli* cells (New England Biolabs) using a published protocol (60), stored in gel filtration buffer supplemented with 5% v/v glycerol, snap-frozen in liquid nitrogen and stored at −80°C until use.

### Generation of CPCs

Purified protein components were reacted together in the sequence indicated in Fig. S1 using published protocols (42, 43, 45). To make the proteinaceous CPC, an excess of SpyCatcher-SnoopCatcher was reacted with SnoopTag-HaloTag-SnoopTag in gel filtration buffer overnight at 4°C with gentle shaking. The covalently conjugated three-part complex was separated from excess starting reagents by gel filtration chromatography (Superdex 200 Increase column) and then reacted with an excess of Capping Protein-SpyTag under the same conditions. A final round of gel filtration chromatography separated the five-part covalent complex from starting reagents.

To prepare half-CPC-DNAs 1 and 2, an excess of SpyCatcher-HaloTag-PCV2 was reacted with Capping Protein-SpyTag, again overnight at 4°C with gentle shaking. The resultant two-part covalent complex was separated from excess starting reagents by gel filtration chromatography, divided in half, and then each half was reacted with an excess of DNA 1 (AAGTATTACCAGAAAAAGGGCCACGGTGGGCCTTGTTTCA) or DNA 2 (AAGTATTACCAGAAACCACACTCCGGTGGACCGAAGCGCGTGAAACAAGGCCCACCGTGGCCCTT) overnight at 4°C with gentle shaking in HUH reaction buffer (50 mM HEPES pH 8.0, 50 mM NaCl, 1 mM MgCl_2_, 1 mM MnCl_2_). The resultant DNA-protein conjugates were separated from excess starting reagents by gel filtration chromatography (Superdex 200 Increase column) and anion exchange chromatography (HiTrapQ column).

### Labeling of ABPs

For TIRF microscopy assays, α-catenins and CPC were thawed and incubated with JF-646-Halo (Promega) for 2 hours on ice in a 1:1 molar ratio, followed by desalting through a Zeba Spin Desalting column (Thermo Fisher) to remove unreacted dye. Labelled proteins were then clarified by ultracentrifugation at 96,460 g in a TLA100 rotor for 20 min at 4°C. Half-DNA-CPC 1 was labeled according to the same protocol, while half-DNA-CPC 1 was labelled with JF-549-Halo (Promega) in the same way, to create a two color CPC-DNA.

### Synthesis of Fibrinogen-GBP

Fibrinogen and GBP were reacted together using published protocols (49). Briefly, fibrinogen (Millipore Sigma) was resuspended in fibrinogen buffer (100 mM NaHCO_3_ pH 8.3, 0.5 mM EDTA) and reacted with an excess of maleimide-PEG8-succinimidyl ester for 1 hour at room temperature, desalted through a Zeba Spin Desalting column (Thermo Fisher), and reacted with an excess of GBP3xCys (prepared according to the same protocol) and mixed overnight at 4°C. Excess free cysteine was added to quench unreacted maleimide groups, and fibrinogen conjugates were precipitated by ammonium sulfate saturation, resuspended in fibrinogen buffer and ultracentrifuged to removed aggregates. Fibrinogen-GBP was snap frozen in liquid nitrogen and stored at −80°C until use.

### Pelleting Assays

Pelleting assays were conducted following the protocol described by Mei et al. (31). Briefly, F-actin was polymerized at 1 μM in G-Mg and KMEI for 1 hour at room temperature. Clarified CPC (20 nM), FL α-catenin or α-cateninΔC (both 1 μM) was added and incubated for 15 min at room temperature. This mixture was then spun down in a TLA-100 rotor (Beckman Coulter) at 9,892 g for 30 min at 4°C to sediment actin filament bundles into a low-speed pellet. The supernatant was subsequently collected and spun down again in a new centrifuge tube in the same rotor at 386,400 g for 30 min at 4°C to sediment single actin filaments into a high-speed pellet. The resultant supernatant was collected, and the low-speed and high-speed pellets were gently washed once with the G-Mg-KMEI solution before resuspension in SDS-PAGE loading buffer. For PAGE analysis, the supernatant and pellets were then run on a 12% NuPAGE Bis-Tris protein gel (Thermo Fisher) followed by Coomassie R250 staining. For the CPC end-binding assay, only a high-speed centrifugation step was conducted, 2 mg/ml BSA was included with F-actin to block centrifuge tubes, and specified F-actin samples were vortexed for 30 s at maximum speed prior to addition of CPC to shear filaments and generate free ends for binding.

For western blot analysis (CPC end-binding assay, Fig. S2A), the supernatant and pellets were run on a 12%Tris-Glycine protein gel (Thermo Fisher) and transferred to a nitrocellulose membrane. The membrane was blocked overnight in blocking buffer (5% DIFCO skimmed milk, 20 mM Tris pH 7.6, 150 mM NaCl, 0.1% Tween20), followed by incubation with anti-HaloTag mAb (Promega) at a 1:2000 dilution in blocking buffer. The membrane was washed 3 times for 5 min in TBST (20 mM Tris pH 7.6, 150 mM NaCl, 0.1% Tween20) before incubation with anti-mouse-HRP (Cell Signaling Technologies) at a 1:2000 dilution in blocking buffer, followed by 3 further washes. Finally, imaging was conducted using western bright ECL buffer (Advansta) on photographic film (Carestream) in a dark room.

### Coverslip preparation

Glass coverslips (Corning 24-60mm, #1.5) were cleaned by sonication in 100% acetone for 30 min and 100% ethanol for 10 min, rinsed with water 3 times, then cleaned by sonication in 2% Hellmanex III liquid cleaning concentrate (HellmaAnalytics) and finally rinsed in water 3 times before air dying. Coverslips were then protein micropatterned (see below) or PEG-silanated by overnight incubation with 1 mg/ml mPEG-silane (Laysan Bio) dissolved in ethanol. Silanated coverslips were washed in water 3 times before air drying and storage at 4°C prior to use.

### Protein micropatterning

The patterning procedure is outlined in Fig. S4. Cleaned coverslips were plasma activated for 5 min in air in an expanded plasma cleaner (Harrick Plasma), and PDMS stencils with 3 mm x 3 mm wells were applied (Alvéole labs). Wells were passivated with 0.1 mg/ml PLL-PEG solution in patterning buffer (10 mM HEPES pH 7.4) for 1-2 hours, then rinsed with DPBS (Gibco) 4 times. Wells were filled with 5 μl PLPP solution (Alvéole labs) and photopatterned using a PRIMO system (Alvéole labs) with UV radiation focused on the glass-buffer interface. For initial characterization of reconstituted networks (Fig. 2), patterns consisted of 10 μm stripes with 10 μm spacing were generated at a dosage of 1000 mJ/cm^2^. For α-catenin experiments (Fig. 3-5), patterns consisted of 2.5 μm stripes with 17.5 μm spacing were generated at a dosage of 2000 mJ/cm^2^, which we empirically found to be more suitable for imaging due to fluorescent background produced by α-catenin’s nonspecific adsorption to the stripes.

Wells were washed 6 times with DPBS and treated according to a modified protocol based on Watson et al. (49). Wells were incubated with 0.1 mg/ml fibrinogen-GBP in patterning buffer for 10-20 min in a humidified petri dish at room temperature. Wells were washed with DPBS 6 times and incubated with 0.5 mg/ml PLL-PEG in patterning buffer for a further 10-20 min. Wells were washed with DPBS 6 times and incubated with 1 mg/ml casein (Sigma) in patterning buffer for a further 10-20 min. Wells were washed with DPBS 10 times and left in DPBS in a humidified petri dish at 4°C until use. At no point was the patterned well allowed to dry out; all wash steps were carried out by removing as much liquid as possible without drying the surface, followed by adding back 10 μl DPBS on top and mixing.

### TIRF force reconstitution assays

Dual-color or tri-color TIRF image sequences (movies) were recorded at room temperature (approximately 25°C) using a Nikon TiE inverted microscope equipped with an H-TIRF module and an Agilent laser launch, driven by Nikon Elements software. Images were taken every 2 s with an Apo TIRF 60X or 100X 1.49 NA objective (Nikon) on an Andor iXon EMCCD camera. 488-phalloidin, 555-phalloidin plus / JF549 and JF646 fluorophores were excited by laser lines at 488 nm, 561 nm and 640 nm, respectively.

For ATP-only assays on unpatterned cover slips (used in PFC and CPC characterization), a PDMS gasket was applied to silanated coverslips to create wells. 1 μM F-actin was pre-incubated with 1, 2, 5 or 10 nM CPC and left for 15 min at room temperature. Each well was prepared immediately before imaging by incubating with 50 nM myosin-5 for 3 min, washed with MB (20 mM MOPS pH 7.4, 5 mM MgCl_2_, 0.1 mM EGTA, 50 mM KCl, 1 mM DTT), followed by incubation with the F-actin / CPC mixture for 30 s, followed by a final wash with MB. An imaging buffer with an oxygen scavenger system and supplemented with calmodulin (MB O.S. 1X ATP: 20 mM MOPS pH 7.4, 5 mM MgCl_2_, 0.1 mM EGTA, 50 mM KCl, 50 mM DTT, 15 mM glucose, 2 mM ATP) was added, and imaging commenced.

For assays on patterned coverslips, 1 μM F-actin was pre-incubated with 1 or 2 nM CPC and left for 15 min at room temperature, then each well was prepared immediately before imaging by first removing the DPBS. Wells were then incubated with 50 nM myosin-5 for 3 min, washed with MB, followed by incubation with the F-actin / CPC mixture for 30 s, then a final MB wash. Imaging buffer was added, and imaging commenced.

For α-catenin experiments a similar procedure was used except that 5 μM FL α-catenin or α-cateninΔC were preincubated with imaging buffer or an identical buffer without ATP (MB O.S. no ATP: 20 mM MOPS pH 7.4, 5 mM MgCl_2_, 0.1 mM EGTA, 50 mM KCl, 50 mM DTT, 15 mM glucose). In paired +/− ATP experiments, the −ATP buffer (including α-catenin) was added and the well imaged for 2 min before recording was paused and two equivalents of +ATP buffer (including α-catenin) were added to the well. Recording resumed immediately (less than 10 s of non-recording). In ATP / apyrase experiments, for the apyrase condition, myosin-5 was incubated with apyrase (New England Biolabs) at 25 units / mL working concentration at room temperature for 10 min before being added to the well, and only −ATP buffer preincubated with α-catenin was used. For the ATP condition, untreated myosin-5 and only +ATP buffer preincubated with α-catenin were used.

### Image analysis and quantification

Image analysis was conducted with FIJI (61), as well as custom python scripts using the sciki-image (62) and NumPy (63) libraries. On unpatterned surfaces, individual PFCs and single filament complexes were manually identified at frame 10 of TIRF microscopy movies and counted to quantify absolute number of filament complexes and proportion of PFCs. Time to rupture was manually recorded for a subset of PFCs, defined as the time between frame 10 and final frame where the intact PFC was observed. For a subset of PFCs, in the frames preceding PFC rupture F-actin fluorescence was recorded in a 21 x 6 pixel box, and CPC fluorescence in a 14 x 14 pixel box. Barbed end concentration was estimated by tracing filaments in experiments in the absence of CPC to measure their average length, estimating the number of subunits expected in a filament of that average length based on the rise of 2.73 nm between subunits on adjacent strands (64), then dividing the G-actin concentration used in the polymerization reaction by that number.

On patterned surfaces, star and Y configuration actin networks were manually identified at frame 10 of TIRF microscopy movies, and time to rupture was recorded, defined as the time between frame 10 and the final frame where network F-actin is seen bridging adjacent stripes. The number of actin bundles seen traversing single 136 μm long stripes was measured by taking a line scan, recording actin fluorescence intensity, and counting the number of peaks.

For α-catenin experiments +/− ATP on patterned surfaces, for each stripe, F-actin fluorescence intensity was threshholded to generate a binary mask, skeletonized, and then dilated to form consistent 5-pixel wide masks containing filament bundle segments. F-actin and α-catenin fluorescence intensity was recorded for each stripe and exemplar heatmaps of the fluorescence intensity ratio in masked areas were generated. For each stripe, the average F-actin fluorescence intensity and average α-catenin : F-actin fluorescence intensity ratio was calculated in the masked regions before and after addition of ATP, then compared.

Four regions of interest featuring interconnected groups of F-actin bundles were identified, grouped into time periods of stable F-actin configurations and averaged to capture different states of the networks. For each average frame, line scans were taken across individual F-actin bundle segments and the average F-actin fluorescence intensity and α-catenin fluorescence intensity of each segment was recorded for calculating the α-catenin : F-actin fluorescence intensity ratio. In one exemplar case (Fig. 4A), masks covering the F-actin bundle segments were generated manually and the underlying α-catenin : F-actin fluorescence intensity ratios were calculated on a per-pixel basis to generate a heat map.

As outlined in Fig. S8, in α-catenin ATP / apyrase experiments, for each stripe, the F-actin fluorescence intensity was thresholded to generate a binary mask, skeletonized, and convolved with a 3 x 3 pixel branch point finding filter to separate individual filament bundle segments. The debranched skeleton was then dilated to form consistent 5-pixel wide masks containing actin filament bundles. The average F-actin intensity and α-catenin fluorescence intensity was measured for each bundle segment to calculate the α-catenin : F-actin fluorescence ratio.

### Plotting and statistical analyses

All plotting and statistical analyses were performed with GraphPad Prism.

## Supplemental Figures

**Figure S1.**
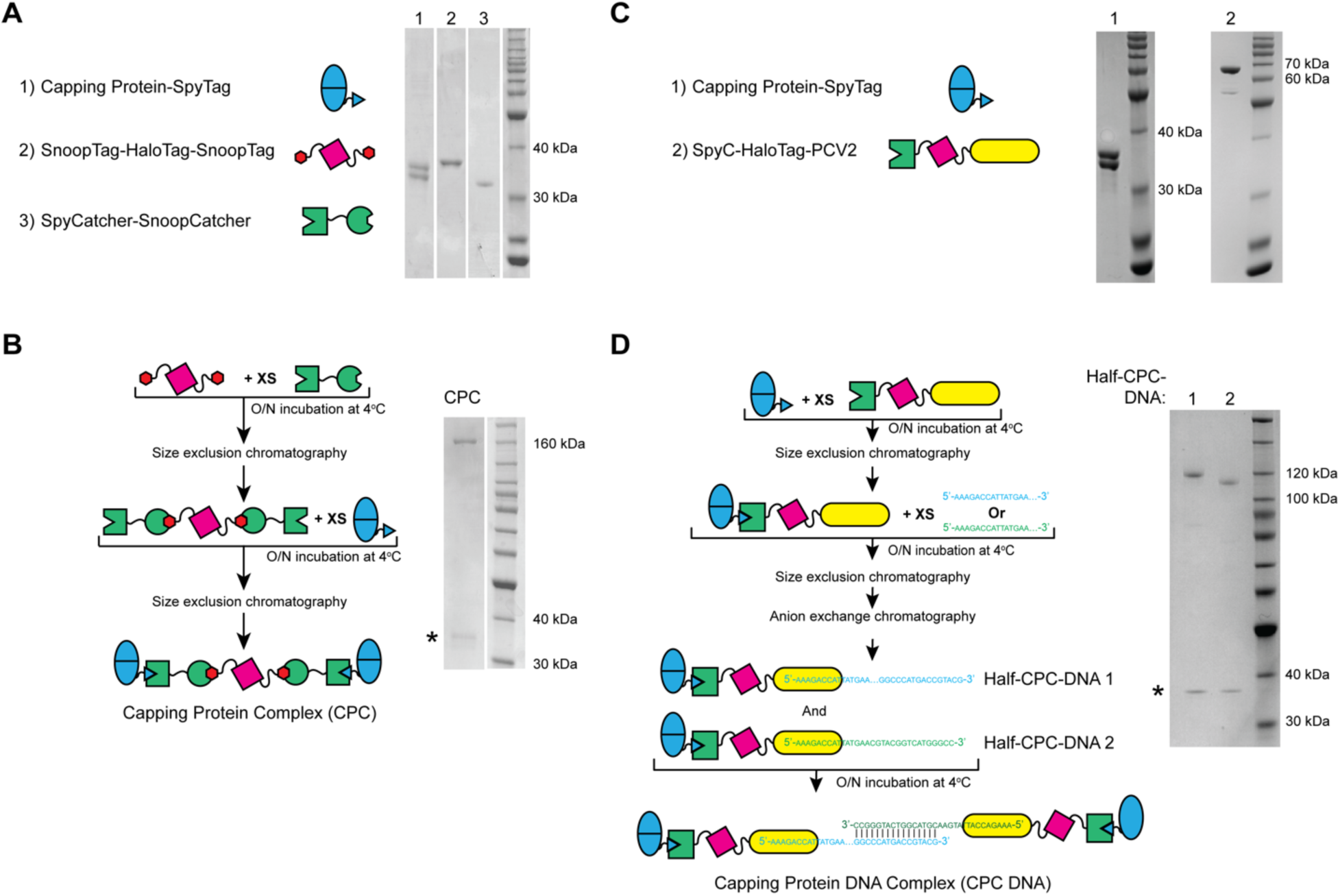
Assembly and purification of CPCs. **A)** SDS-PAGE analysis of proteinaceous CPC components. **B)** Reaction scheme and SDS-PAGE analysis of the assembled five-part CPC. Asterisk indicates CapZA, which is non-covalently associated with the CPC. **C)** SDS-PAGE analysis of CPC-DNA components. **D)** Reaction scheme and SDS-PAGE analysis of the two halves of the assembled CPC-DNA. Asterisk indicates CapZA.

**Figure S2.**
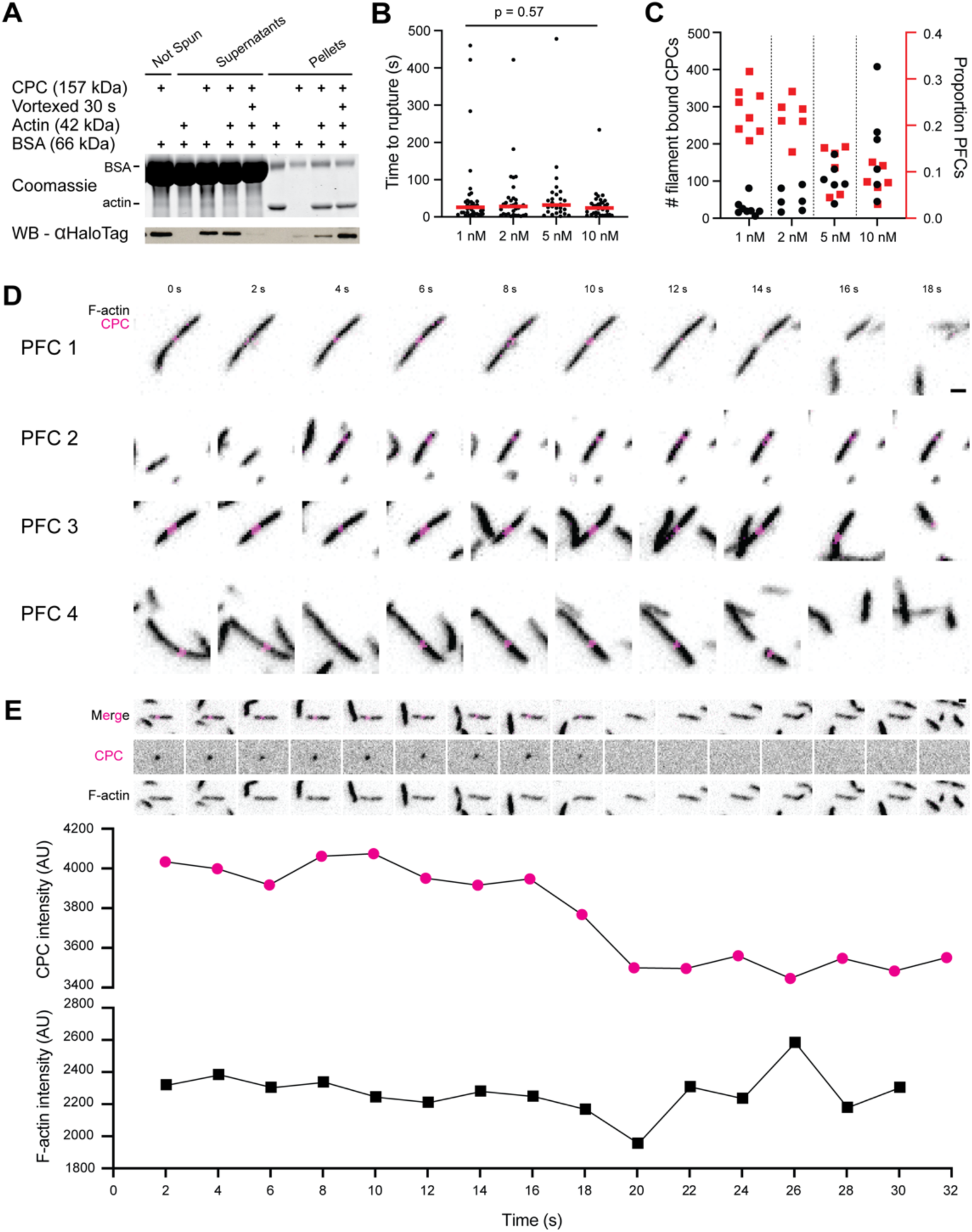
Characterization of PFCs assembled with CPC. **A)** SDS-PAGE analysis of CPC / F-actin co-sedimentation assay. 2 mg / ml BSA was included as blocking reagent. Vortexing filaments shears them, increasing the free barbed-end concentration. Enhanced CPC co-sedimentation under this condition indicates end-dependent binding. **B)** Time to rupture for PFCs from Fig. 1C, pooled by CPC concentration. Bars indicate means. n = 38 (1 nM); 33 (2 nM); 27 (5 nM); 32 (10 nM) from 26 independent trials. KW test with Dunn’s correction for multiple comparisons. **C)** Replotting of data from Fig. 1D, showing all data points for number of CPCs bound to filaments (black) and proportion of CPCs engaged in PFCs (red) versus CPC concentration. n = 7 (1 nM); 6 (2 nM); 6 (5 nM); 6 (10 nM) independent trials. **D)** Additional montages of PFCs breaking under tension over time. Scale bar, 1 μm. **E)** Top: montage where CPC bleaches prior to PFC rupture. Scale bar, 1 μm. Bottom: fluorescence intensity of F-actin and CPC in each frame. One-step bleaching of CPC indicates the presence of a single molecule.

**Figure S3.**
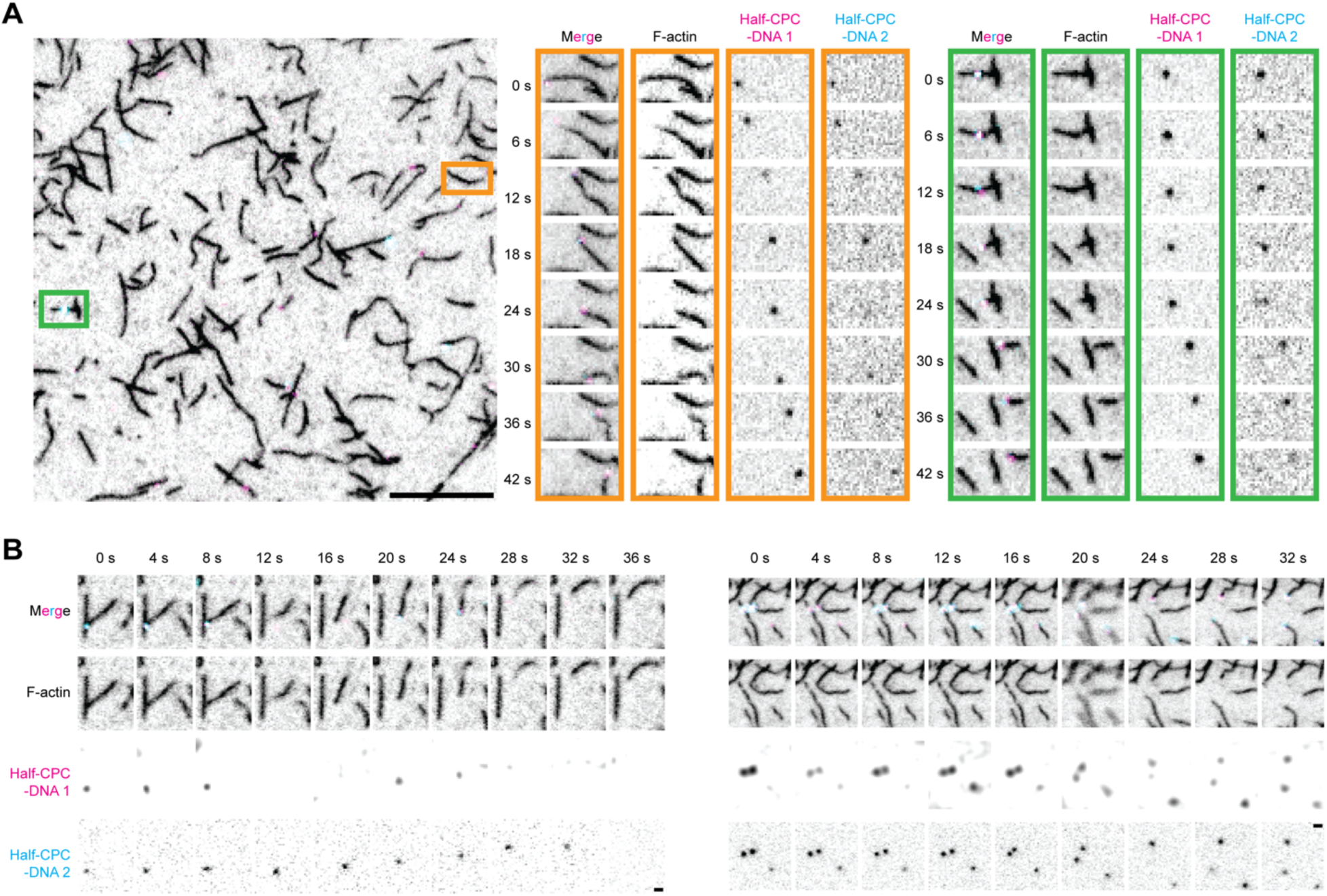
Characterization of dPFCs assembled with CPC-DNA. **A)** Left: micrograph of a field of dPFCs prepared with CPC-DNA. Scale bar, 10 μm. Right: montage of single filament with a complete CPC-DNA attached moving over time (orange) and montage of dPFC breaking over time (green). **B)** Montages of additional dPFCs breaking under tension over time. Scale bar, 1 mm.

**Figure S4.**
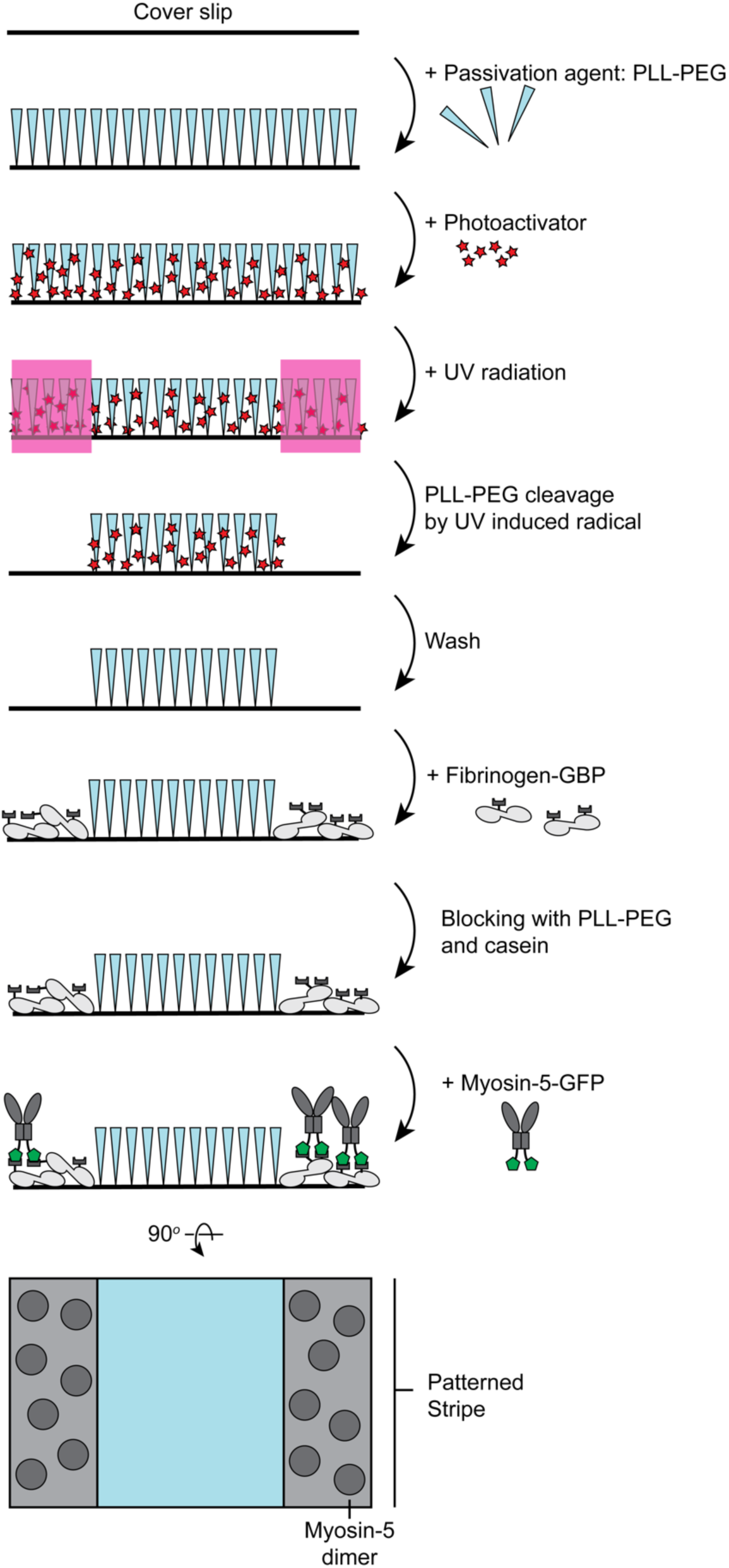
Protein micropatterning procedure. Cartoon of PRIMO UV photopatterning procedure showing two adjacent stripes of myosin-5 on a coverslip (see Methods for details).

**Figure S5.**
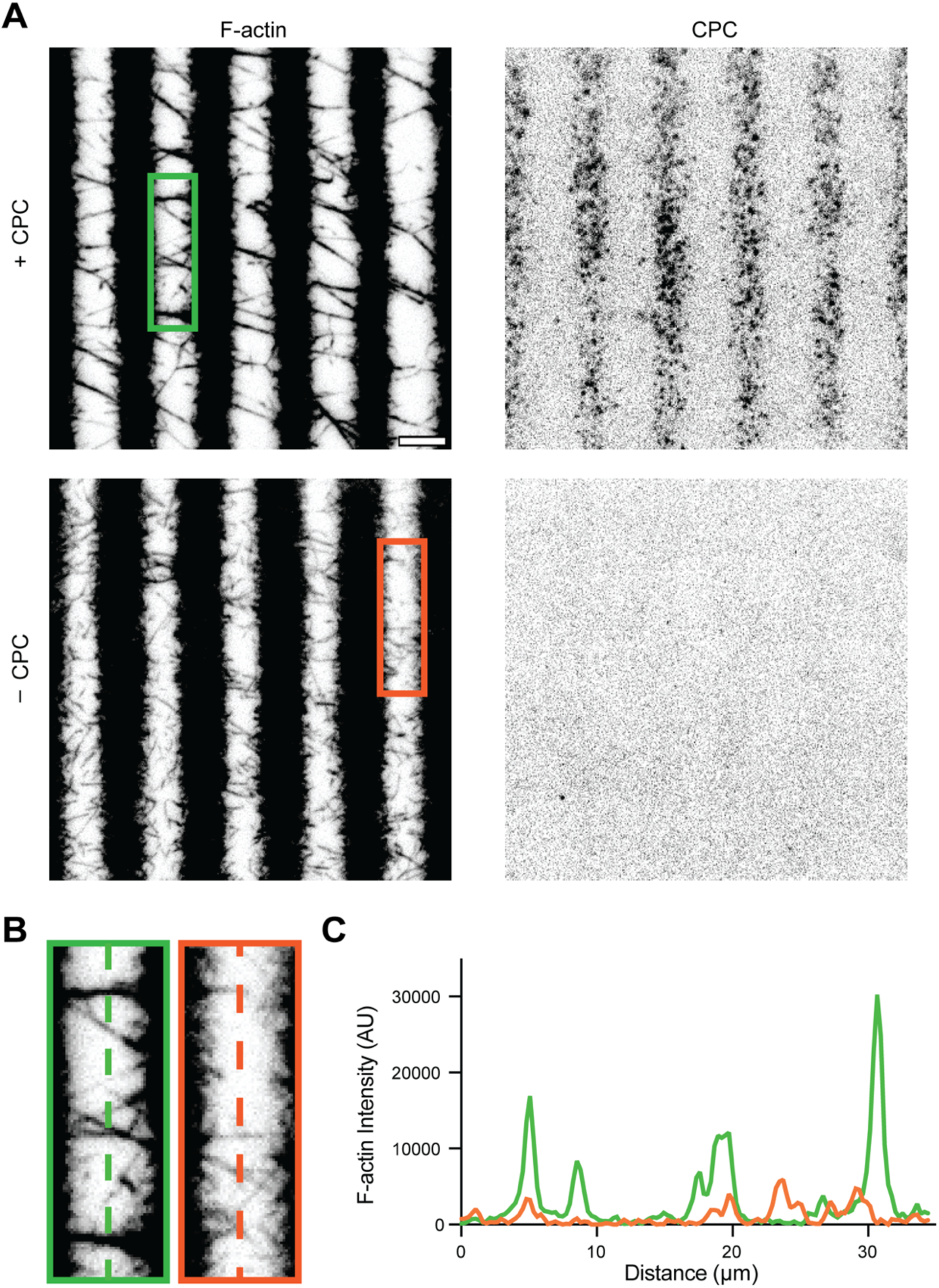
“Star” and “Y” configuration F-actin networks form in the presence of CPC. **A)** Micrographs of F-actin on micropatterned fields of myosin-5 in the presence (top) and absence (bottom) of CPC. Scale bar, 10 μm. **B)** Detail views of gaps between stripes in both conditions selected for linescan analysis (dotted lines). **C)** F-actin intensity profile along linescans: + CPC condition, green; - CPC condition, orange. The lower intensity of peaks in the - CPC condition indicates reduced F-actin bundling.

**Figure S6.**
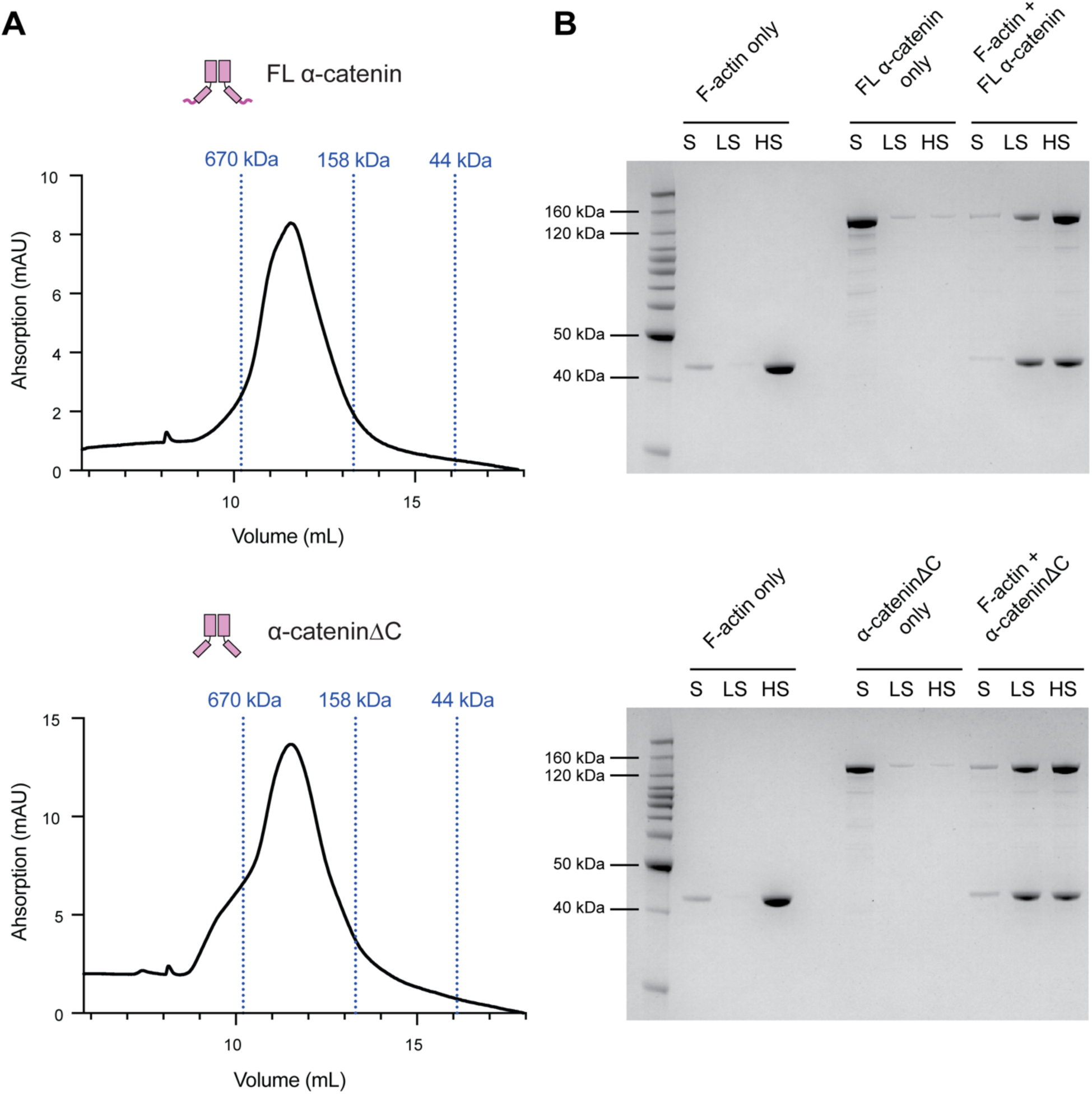
Purification and characterization of FL α-catenin and α-cateninΔC. **A)** Size exclusion chromatography profiles of purified FL α-catenin (top) and α-cateninΔC (bottom). Molecular weights of Halo-tagged FL α-catenin and α-cateninΔC are 128 kDa and 124 kDa, respectively. Dotted lines indicate retention volumes of specified molecular weight standards. **B)** SDS-PAGE analysis of co-sedimentation assays of FL αCatenin (top) and αCateninΔC (bottom). S = supernatant, LS = low speed pellet, HS = high speed pellet.

**Figure S7.**
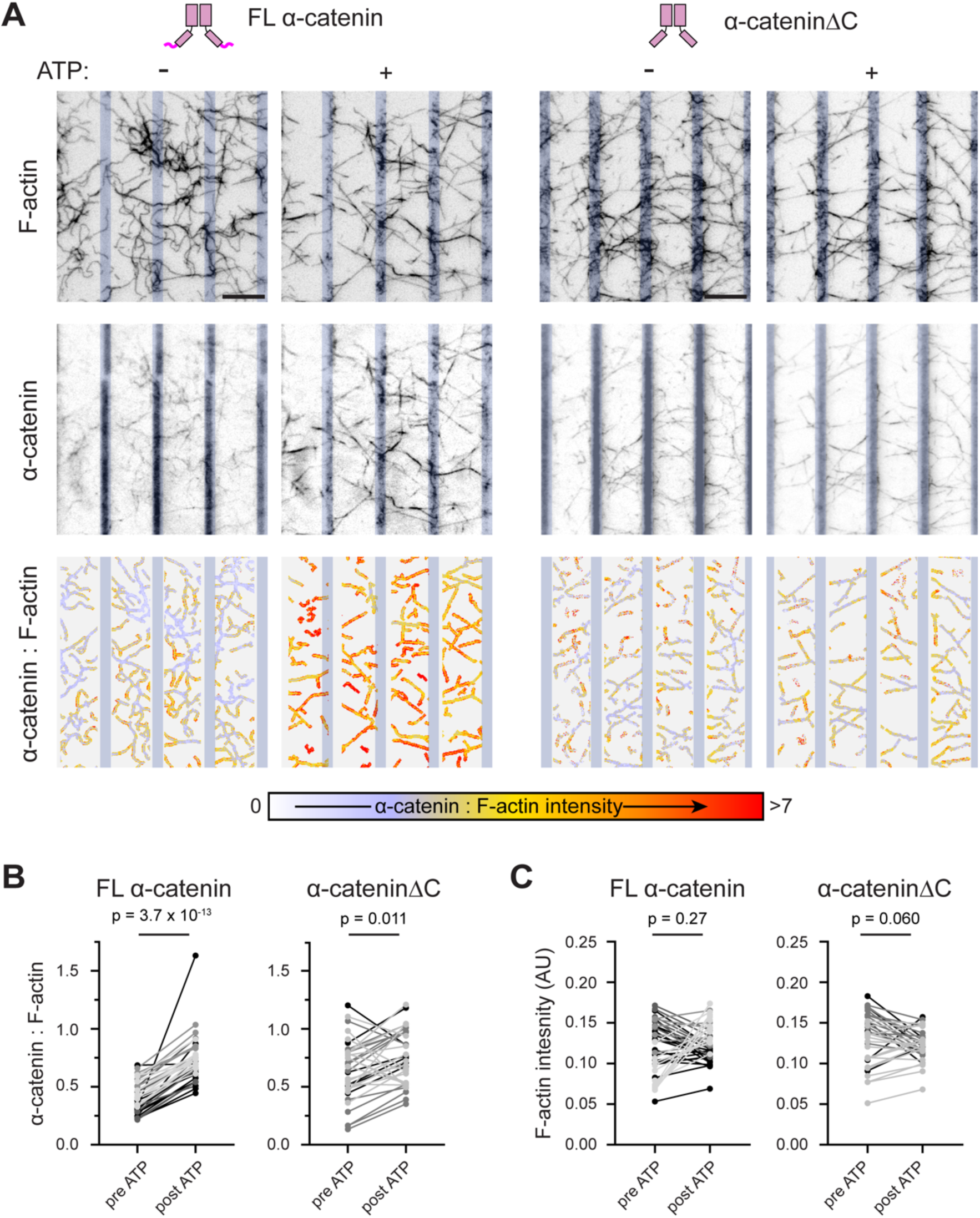
Dimeric FL α-catenin features force-activated F-actin binding activity. **A)** Top: micrographs of PFC networks in the presence of FL α-catenin (left) or α-cateninΔC (right) before and after the addition of ATP. Bottom: α-catenin : F-actin fluorescence intensity ratio images of the same fields of view. Vertical bars indicate positions of myson-5 stripes. Scale bar, 10 μm. **B)** Paired analysis of average α-catenin : F-actin fluorescence intensity ratio per inter-stripe gap before and after ATP addition for FL α-catenin (left) or α-cateninΔC (right). Welch’s t-test: n = 40 gaps from 4 independent trials (FL α-catenin); 35 gaps from 3 independent trials (ΔC). Shades of grey indicate points from different trials. **C)** Paired analysis (Welch’s t-test) of actin fluorescence intensity from same data presented in **B**.

**Figure S8.**
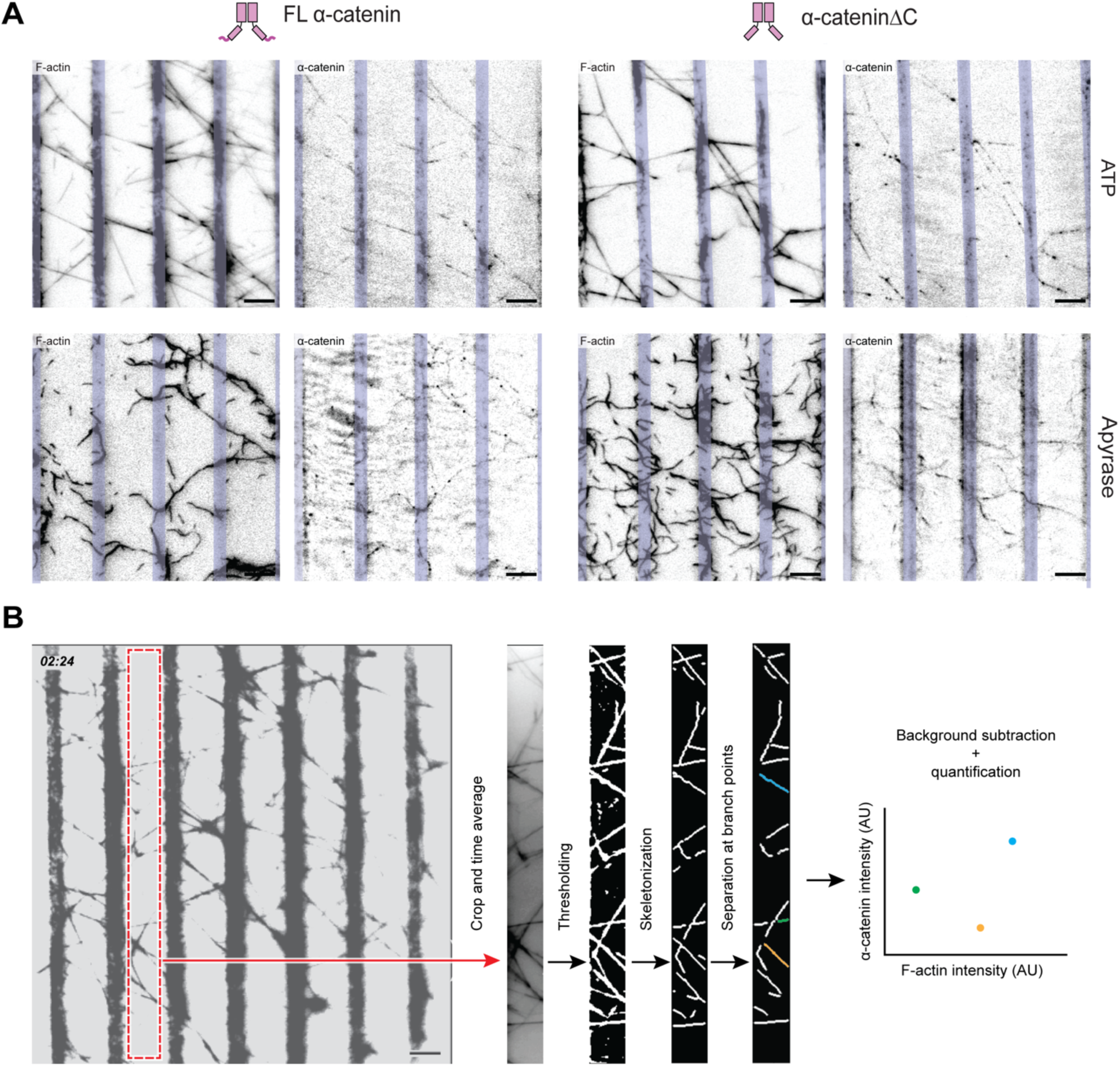
Image segmentation and quantification pipeline. **A)** Representative micrographs of PFC networks in the presence of either FL α-catenin (left) or α-cateninΔC (right) treated with either ATP (top row) or apyrase (bottom row). Fields of view are identical to those depicted in the α-catenin : F-actin fluorescence intensity ratio images in Fig 4A. Vertical bars indicate the positions of patterned myosin-5 stripes. Scale bar, 10 μm. **B)** Cartoon showing how bundle segments are identified and segmented from networks for analysis. See Methods for details. Scale bar, 10 μm.

## Movie Legends

**Movie S1: PFCs prepared with CPC come under tension on a random field of myosin-5 motors.** Same field of view as Fig. 1B. F-actin is black; CPC is magenta. Green arrows highlight PFCs, which turn red immediately before PFC rupture. Scale bar, 10 μm.

**Movie S2: Half-CPC-DNA 1 translocates on barbed ends of actin filaments in a gliding assay.** From a single field of view, left frame displays half-CPC-DNA 1 (magenta), right frame displays F-actin (black). In addition to mobile half-CPC-DNAs, some molecules adhere to the surface and do not translocate. Scale bar, 10 μm.

**Movie S3: Half-CPC-DNA 2 translocates on barbed ends of actin filaments in a gliding assay.** From a single field of view, left frame displays half-CPC-DNA 2 (cyan), right frame displays F-actin (black). The sample features a mixture of mobile and immobile half-CPC-DNAs. Scale bar, 10 μm.

**Movie S4: dPFCs made with CPC-DNA under tension on a random field of myosin-5 motors.** Same field of view as Fig. 1E. F-actin is black; half-CPC-DNA 1 is magenta; half-CPC-DNA 2 is cyan. Green arrows highlight dPFCs, which turn red immediately before dPFC rupture. Scale bar, 10 μm.

**Movie S5: higher-order PFC networks under tension on micropatterned stripes of myosin-5.** Same field of view as Fig. 2A. F-actin is black, CPC not shown. Regions featuring uniform high density of F-actin correspond to myosin-5 stripes. Scale bar, 10 μm.

**Movie S6: FL α-catenin accumulates on PFC networks in response to tension.** From a single field of view, left frame displays F-actin, right frame displays FL α-catenin. Recording was initiated after ATP addition to activate motors. Regions featuring uniform high density of F-actin and α-catenin correspond to myosin-5 stripes. Scale bar, 10 μm.

**Movie S7: FL α-catenin dynamically localizes within a mechanically rearranging PFC network.** From a single field of view of network analyzed in Fig. 3, left frame displays F-actin, middle frame displays FL α-catenin. Right frame displays time-averaged F-actin signal between substantial transitions assigned as states. Patterned stripes are not visible but are present to the immediate left and right of the frame. Scale bar, 2 μm.

